# DeepDynamics resolves cell-subtype and clinicopathological dynamics from bulk RNA-seq to identify mediators of Alzheimer’s disease risk

**DOI:** 10.64898/2025.12.15.694345

**Authors:** Yifat Haddad, Yuval Rom, Gilad Sahar Green, Anael Cain, Barak Raveh, Naomi Habib

## Abstract

Processes such as Alzheimer’s disease and aging are shaped by dynamic cascades of cellular and molecular changes. Mapping how genetic and environmental risk factors alter these cascades is critical for pinpointing disease susceptibility and guiding therapeutic intervention, yet this task requires dense sampling of cell-subtype and -subpopulation changes throughout the full course of disease. Bulk measurements provide scale but obscure cell-type and subpopulation resolution, whereas single-cell assays offer resolution but lack sufficient sample size and temporal coverage. We present DeepDynamics, a deep-learning framework that combines a smaller reference sc/snRNA-seq atlas with a larger bulk RNA-seq cohorts to infer cell-subpopulation and clinicopathological dynamics along annotated trajectories. Applied to Alzheimer’s disease (AD), DeepDynamics mapped 1,092 cortical bulk profiles onto aging and disease trajectories, revealing faster accumulation of AD clinicopathologies, disease-associated glia, and specific neuronal subpopulations in *APOE4* carriers. These changes, stronger in females, were accompanied by upregulation of cell-type-specific molecular pathways and early downregulation of heat-shock proteins across cell types. By unifying genetic risk, pathology, and cell-subpopulation dynamics, DeepDynamics provides a scalable approach to prioritize putative mediators of risk and resilience amid the myriad changes along the cascade, thus informing targeted therapies.

## Introduction

Multifactorial biological processes such as Alzheimer’s disease (AD) and aging unfold through dynamic cascades of cellular and molecular changes, themselves shaped by genetic and environmental risk factors^1–3^. In AD, the *APOE ε4* (*APOE4*) allele increases disease risk by three- to fourfold compared with the neutral variant *APOE ε3* (*APOE3*)^4^, and females face a ∼twofold higher risk than males^5^. However, the cell types and molecular pathways mediating these risk factors and others remain largely unresolved. Linking risk factors such as *APOE4* and sex to underlying cellular and molecular mechanisms across disease stages is crucial for differentiating between protective and harmful processes and for prioritizing therapeutic targets among the many observed cellular and molecular changes (e.g.^6–11)^. Addressing this task requires continuous profiling of cell-subpopulation and clinicopathological dynamics along disease progression within large, diverse cohorts, to provide the temporal coverage, sample size, and resolution.

Bulk tissue RNA-seq can be used to profile large and diverse cohorts (e.g.^12,13^), but since transcript levels are averaged across all cells in the sample, it lacks cell-type and subpopulation resolution, hindering the mapping of molecular changes to specific cell subpopulations. By contrast, single-cell and single-nucleus RNA-sequencing (sc/snRNA-seq) enable high-resolution mapping of cellular heterogeneity across conditions^14–22^. Specifically, such large-scale snRNA-seq profiling of human aging and AD brains^23,24^ identified a coordinated shift to disease-associated states across glial cell types, as well as vulnerability of specific neuronal subtypes^15–18^.

Trajectory inference algorithms extend transcriptomic data analyses from static snapshots to alignment of cells along a continuous pseudo-timeline (*e.g.* ^25–28)^. Similar inference methods can also be applied at the subject-level to order human samples along a pseudo-timeline of a given process based on sc/snRNA-seq measurements (*e.g.*^23,29^). Since each sample captures only a single unknown time point along disease or aging, such ordering is critical for downstream analyses: it separates early from late disease stages, untangles overlapping processes, and enables reconstruction of the underlying continuous cell-subpopulation and clinicopathological dynamics. Such subject-level trajectory inference was used to align hundreds of snRNA-seq-profiled aging human brains along two distinct trajectories disentangling the overlapping processes of AD progression and alternative brain aging, revealing diverging cellular dynamics along each one^23^, including a coordinated shift toward disease-associated glial cell subpopulations^15–18^ in the progression of AD (prAD) trajectory^23^.

However, the scale of sc/snRNA-seq datasets remains limited due to cost and labor, hindering stratified analysis by genetic and clinical subgroups. The need for both scale and resolution becomes acute when considering cell-subpopulation dynamics, which requires dense sampling from each subgroup throughout the full course of disease. A promising solution for addressing the limited size of single-cell datasets is to integrate large-scale bulk-profiled cohorts with smaller higher-resolution single-cell datasets, an approach validated for inferring static cellular compositions^24,30–32^, yet methods for inference of cell-type and subpopulation dynamics directly from bulk data have not been established.

To address this gap, we developed **DeepDynamics**, a framework integrating the scale of bulk RNA datasets with the resolution of single-cell data, by reconstructing cell-subpopulation dynamics along trajectories of biological processes from the bulk datasets. DeepDynamics trains a deep neural network (DNN) on an annotated sc/snRNA-seq reference atlas and overlapping bulk samples, followed by applying the DNN over the larger bulk RNA cohort. It employs explainability analysis of the DNN^33^ to identify cellular subpopulations predictive of the progression of the process, then comparatively analyzes the cellular dynamics of these subpopulations across genetic or clinical groups, benefitting from the increased scale of the bulk cohort, thus prioritizing putative cellular drivers and therapeutic targets. We further show that DeepDynamics can be repurposed for flexible integration of additional snRNA-seq samples to enable cell-type-specific molecular investigation at scale. Overall, we present a broadly applicable method that offers proof-of-principle for leveraging bulk datasets and provides a roadmap for future experimental design.

We applied DeepDynamics to investigate the effects of the AD risk factors, *APOE4* and sex, in 1,092 bulk RNA-seq profiles of aging human dorsolateral prefrontal cortex (DLPFC) from the ROSMAP cohort, a longitudinal study of aging providing detailed clinical and pathological information^34,35^. Trained on a smaller fully-annotated snRNA-seq atlas^23^ spanning two trajectories leading to progression of AD (prAD) and alternative brain aging (ABA), DeepDynamics mapped cell-subpopulation and clinicopathological dynamics along each trajectory. We revealed a previously unmapped accelerated accumulation of clinicopathologies in *APOE4* carriers with a sex-dependent effect on tau tangles (tau) and a sex-independent effect on amyloid-β (Aβ). At the cellular level, these effects were mirrored by increased vulnerability of specific neuronal subtypes and accelerated accumulation of disease-associated glia subpopulations, particularly in females. At the molecular level, we identified altered cell-type-specific molecular pathways in *APOE4* carriers, as well as a decrease in expression of heat-shock genes across cell types. Overall, the scale and temporal resolution provided by DeepDynamics revealed *APOE4*-dependent modulations at the pathological, cellular, and molecular levels, and how these are further shaped by its interaction with sex. These findings highlight putative mediators of AD risk and demonstrate the value of stratifying cellular dynamics by genetic risk to elucidate disease mechanisms.

## Results

### DeepDynamics aligns bulk samples along trajectories of aging and disease to investigate cell-subpopulation dynamics

To examine how *APOE4* genotype and sex shape AD risk at the pathological, cellular and molecular level, we developed DeepDynamics (**Fig. 1a-c, Extended Data Fig. 1a,b**). DeepDynamics offers a scalable, statistically robust framework for linking risk factors to cellular and molecular cascades, yielding causal mechanistic insights to prioritize therapeutic targets.

**Figure 1.**
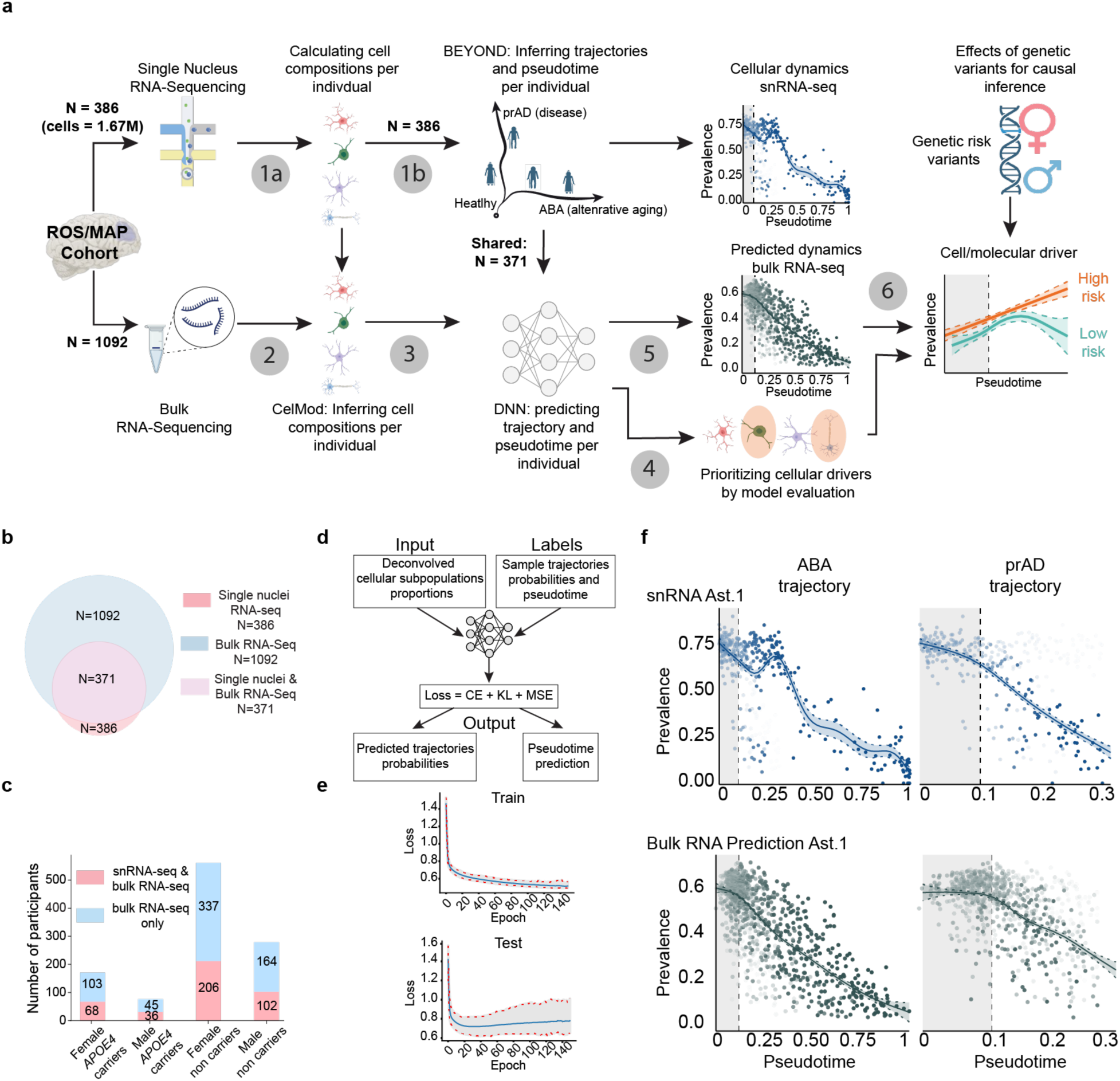
DeepDynamics framework for inferring cell-subpopulation dynamics from bulk RNA-seq data based on pre-annotated single-cell data. **(a)** Overview of the DeepDynamics framework. Starting from an annotated snRNA-seq atlas (step 1a), samples are aligned along inferred trajectories of biological processes (1b). DeepDynamics is trained on matched snRNA-seq and bulk RNA-seq samples to predict cellular compositions (2), and to predict trajectory assignment and pseudotimes across the full bulk cohort using a DNN (3). Model explainability is used to rank putative cellular drivers of the process (4). Dynamics of both clinicopathological and cellular prevalence are modeled along trajectories (5) and compared between conditions (6). The framework was applied to bulk RNA-seq profiles from 1,092 cortical samples, trained on an snRNA-seq atlas of 386 individuals aligned to two trajectories – ABA (alternative brain aging) and prAD (progression of Alzheimer’s disease)^23^ - enabling stratified analysis of AD risk factors *APOE4* genotype and sex. **(b)** Study cohort composition, showing the number of samples with snRNA-seq, bulk RNA-seq and overlapping profiles. **(c)** Improved coverage within the larger bulk cohort. Distribution *of APOE* genotype and sex across the overlapping snRNA-seq/bulk-RNA-seq samples and the additional bulk-only samples. **(d)** Schematic of the DNN model input and output within DeepDynamics. **(e)** Train and test loss-curves of the DNN model across training epochs. **(f)** Consistent dynamics and dense coverage in the bulk inferred cellular dynamics. The prevalence of homeostatic astrocyte subpopulation Ast.1 (y-axis) along pseudotime (x-axis), in the snRNA-seq data (top, *n*=386) and bulk RNA-seq predicted (bottom, *n*=1,092), along the ABA (left) and prAD (right) trajectories. See **Extended Data** Fig 1**-2.**

DeepDynamics pipeline overview (**Fig. 1a**): As input, it is provided with an annotated sc/snRNA-seq atlas with detailed cell subpopulations and a partially overlapping large bulk RNA-seq cohort. In step 1, cellular compositions (*i.e.* the prevalence of each cell subpopulation) are derived per individual in the sc/snRNA-seq cohort and used as input for trajectory inference algorithms (*e.g.* BEYOND^23,25^) that predict trajectories that capture cellular changes along the progression of one or more biological processes (*e.g.* healthy aging *vs.* disease). Each individual is assigned a probability per trajectory and mapped to a pseudotime along it. In step 2, we infer cell-type compositions for each bulk RNA sample (*e.g.* by CelMod deconvolution algorithm shown to accurately detect also cell subpopulations in bulk data^24^). In step 3, predicted trajectory probabilities and pseudotime are assigned to each bulk RNA sample by a deep neural network (DNN). In steps 2 and 3, training is conducted on overlapping sc/snRNA and bulk samples, and applied to the remaining bulk samples. In step 4, informative cellular events that might be affecting the progression along each trajectory are prioritized by model explainability analysis, such as SHapley Additive exPlanations (SHAP)^33^. In step 5, cellular, molecular and pathological dynamics are inferred for the full bulk dataset by Generalized Additive Models^36^ (GAMs). In step 6, dynamics are compared across cohorts, such as high- *vs.* low-risk genotypes, benefiting from the statistical power of the larger bulk dataset.

To study AD risk mechanisms we applied DeepDynamics on a well-annotated snRNA-seq atlas of DLPFC cortical samples defining 95 cell subpopulations from 386 individuals^23^ and a bulk RNA-seq dataset from 1,092 individuals (**Fig. 1b, Supplementary Table 1**), randomly sampled from the ROSMAP cohort^34,35^. Both datasets span cognitively healthy, mildly impaired, and advanced AD, with varying levels of neuropathological hallmarks (**Extended Data Fig. 1c**, **Fig. 1c; Extended Data Fig. 1a, b**). DeepDynamics trained on 371 individuals profiled by both snRNA/bulk (**Fig. 1d,e**), then projected the expanded bulk RNA-seq cohort along two previously-defined trajectories: one capturing progression of AD (prAD) and the other alternative brain aging (ABA)^23^ (**Fig. 1f**). This alignment mapped the temporal ordering of cellular transitions and clinicopathological traits along the progression of AD and their distinction from brain aging in all 1,092 individuals (**Supplementary Table 1-2**). By constraining the analysis to previously-validated trajectories of aging and disease, we could examine how risk factors modulate these processes at the pathological, cellular, and molecular levels.

DeepDynamics accurately predicted trajectory probabilities and pseudotime in the bulk dataset (**Fig. 1e**; **Extended Data Fig. 1d,e**), recapitulating the snRNA-seq measured dynamics along both trajectories in all subpopulations (**Fig. 1f, 2a; Extended Data Fig. 2a**), preserving the directionality and magnitude of observed trends and smoothing the signal (**Extended Data Fig. 2b**). It was also repurposed to flexibly integrate additional non-annotated snRNA-seq samples by aligning them along existing trajectories (**Extended Data Fig. 1f, g**). This expansion, from 371 to 1,092 aligned samples, enhanced statistical power and temporal sampling density, facilitating robust comparisons across subgroups of individuals.

### Explainability analysis reveals potential cellular drivers of AD and aging

We applied SHAP^33^ to interpret the DeepDynamics DNN, ranking the input features (prevalence of cell subpopulations) by their average contribution to the predictions (**Fig. 1a**). The SHAP rankings prioritizes the cell subpopulations that were most informative for predicting the three outputs: prAD probability, ABA probability and pseudotime (**Fig. 2b; Extended Data Fig. 3a**). The rankings were consistent across the full bulk cohort (n=1,092), the overlapping bulk and snRNA-seq (n=371; **Extended Data Fig. 3b**) and the independent bulk-only set (n=721; **Extended Data Fig. 3c**). We interpreted the most informative cell subpopulations for trajectory prediction as *putative cellular drivers* of the process and prioritized them for follow-up analysis (**Fig. 2c-f**).

**Figure 2.**
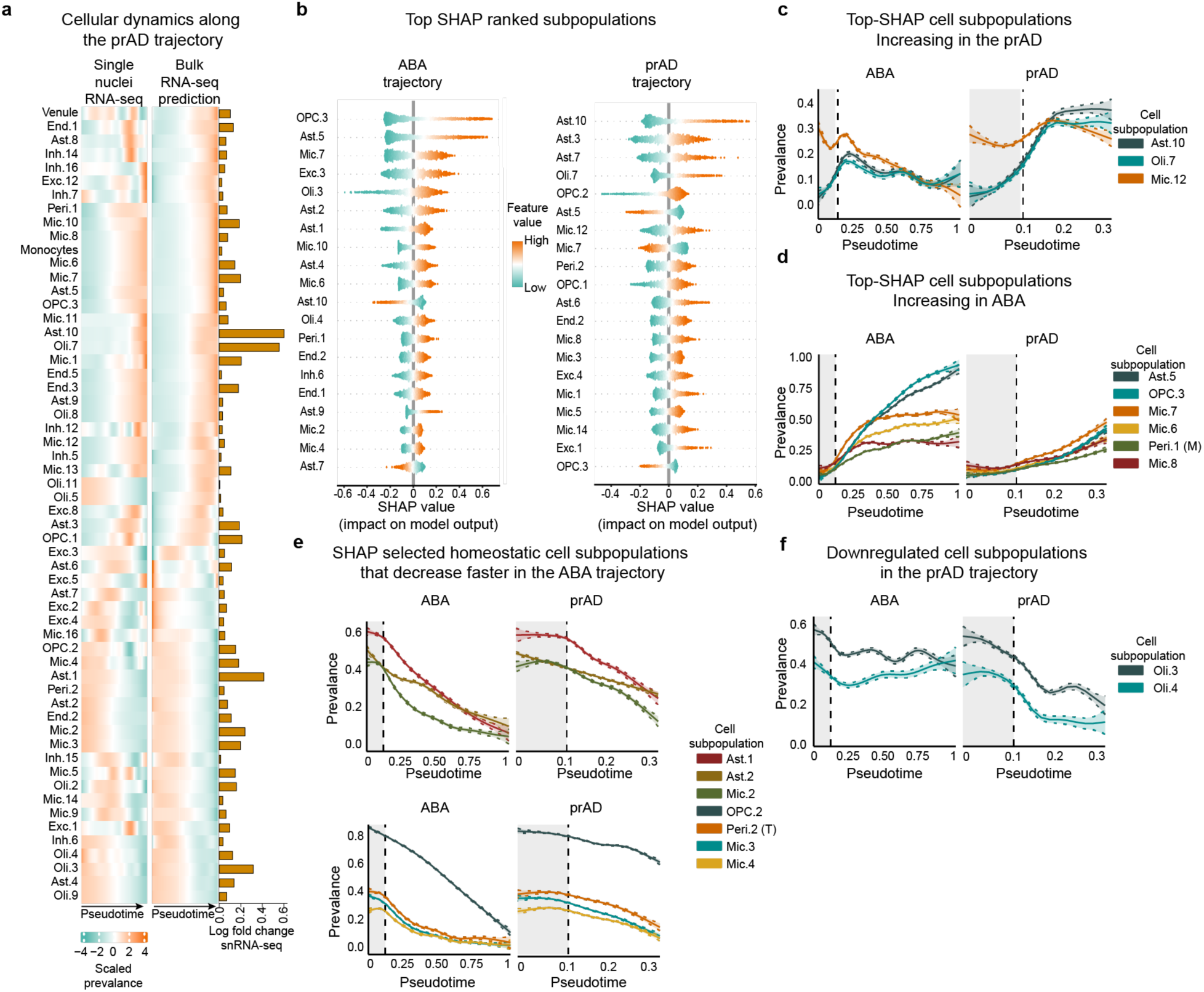
SHAP explainability analysis of the DeepDynamics DNN model identifies putative cellular drivers of AD and aging. **(a)** Bulk-RNA-predicted cellular dynamics match snRNA-seq. Scaled GAM estimates of prevalence (proportion) for each cell subpopulation (rows) as a function of pseudotime along the prAD trajectory (y-axis) for snRNA-seq analyzed by BEYOND^23^ and bulk RNA-seq by DeepDynamics. Horizontal bar plot (right): magnitudes of log-fold change (LFC) between minimum and maximum prevalence values for each subpopulation in the snRNA-seq dataset; Smaller LFC values indicate relatively static dynamics. **(b)** SHAP explainability analysis identifies informative cell subpopulations. Distributions of SHAP values (x-axis) in the bulk dataset for top-ranked cell subpopulations (y-axis), ranked by mean absolute SHAP values quantifying the impact of each subpopulation on predictions for the ABA (left) or the prAD (right). Colors denote feature values (subpopulation prevalence). **(c–f),** SHAP-prioritized cell subpopulations with distinct dynamic patterns: increasing along prAD (**c**), increasing along ABA (**d**), faster decline along ABA vs. prAD (**e**), and downregulated in prAD (**f**). lines = GAM estimates of prevalence of each cell subpopulation along the pseudotime. Dashed lines = confidence intervals. See **Extended Data** Fig 3.

For the prAD trajectory, SHAP prioritized known disease-associated glial cell subpopulations^23^, including the astrocyte subpopulation Ast.10, oligodendrocyte subpopulation Oli.7 and the microglia subpopulation Mic.12, whose prevalence increased with disease (**Fig. 2c**), alongside Oli.3 and Oli.4, whose prevalence declined (**Fig. 2f)**. For the ABA trajectory, Ast.5 and oligodendrocyte precursor cell (OPC) subpopulation OPC.3 ranked highest, consistent with the selective increase in their prevalence along the ABA trajectory (**Fig. 2d**). SHAP also captured subtler patterns, such as the accelerated loss of homeostatic glial subpopulations along the ABA (**Fig. 2e**), and trajectory-specific shifts in rare pericyte subpopulations^37^ (**Fig. 2d,e, Extended Data Fig. 3d**). Thus, DeepDynamics uncovers both known and previously overlooked cell subpopulations that may shape aging and disease cascades.

### *APOE4* and sex are linked to aggravation of AD clinicopathology

By enhancing the sample coverage along the progression of the prAD and ABA trajectories (n=1,092 bulk profiles), DeepDynamics has enabled us to investigate the effects of *APOE* genotype and sex on AD clinicopathology and cellular dynamics. To this end, we utilized detailed quantitative pathological and clinical data within the ROSMAP cohort, including continuous estimates of AD pathology load - Aβ and tau - and of the rate of cognitive decline.

We confirmed the known association of these three AD traits and *APOE* genotype or sex^4,38–40^ within the bulk RNA cohort. Linear regression analysis (confounder-adjusted, FDR<0.01; **Methods**; **Fig. 3a**) showed that *APOE4* carriers (homozygotes or heterozygotes) were associated with higher Aβ and tau loads, as well as faster cognitive decline, compared with non-carriers (including *APOE3* and *APOE2* genotypes). Similarly, females showed higher tau load and an accelerated rate of cognitive decline compared with males, though notably, with no significant sex effect on Aβ load (**Fig. 3b**)^38–40^.

**Figure 3.**
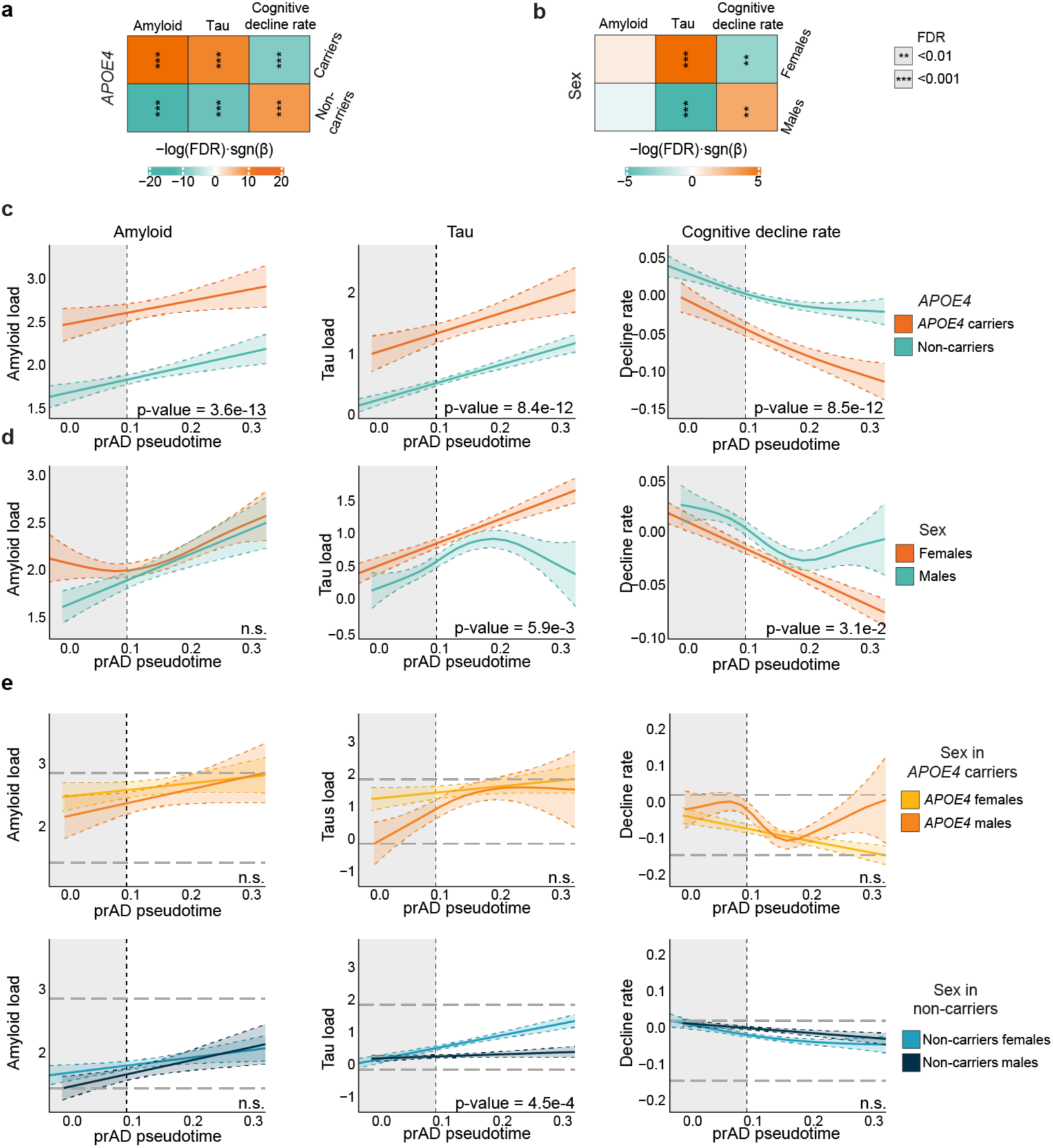
Effect of *APOE4* and sex on clinicopathological dynamics in AD. **(a)** Heatmap of linear-model associations between AD clinicopathologies and *APOE4* in the bulk cohort (*n*=1,092; *β*=effect size; *FDR*=Benjamini-Hochberg-adjusted p-value). **(b)** Same as (**a**) but for associations with sex. **(c-e),** DeepDynamics-inferred AD clinicopathological dynamics along the prAD pseudotime (x-axis, n=1,092), stratified by: *APOE4* genotype (**c**), sex (**d**), or combined stratification by *APOE4* and sex (**e**). P-values were computed using an ANOVA test. n.s. = non-significant. See **Extended Data** Fig 4.

To elucidate how these risk factors affect disease progression, we modeled the dynamics of AD traits along the prAD trajectory, spanning disease stages from cognitively healthy individuals with minimal or no AD-pathologies to advanced AD with clinical dementia and high pathology load^23^. For each stratified group, we fitted non-linear GAMs^36^ and assessed group differences using an ANOVA test for stratified *vs.* pooled models (**Supplementary Table 3**). We found that *APOE4* carriers, compared to non-carriers, showed higher pathology load and faster cognitive decline, emerging in early prAD stages and persisting throughout (p-value<8e-12; **Fig. 3c**). Conversely, compared to males, females showed accelerated tau accumulation (p-value<5.9e-3) and cognitive decline (p-value<3.1e-2), primarily at later disease stages, with no significant difference in Aβ load along pseudotime of the prAD trajectory (**Fig. 3d**).

When examining the *APOE4*-sex interplay, we found no significant interaction between the effect of *APOE4* and sex on Aβ load and the rate of cognitive decline (p-value>0.05, **Fig. 3e**). By contrast, tau dynamics revealed a nuanced interaction between sex and *APOE* genotype. Among non-carriers, females showed a significantly faster rate of tau accumulation, ultimately reaching levels comparable to *APOE4* carriers (p-value<5e-4, **Fig. 3e**). Among *APOE4* carriers, females exhibited higher tau load at earlier disease stages, though the difference between males and females across the entire trajectory was not statistically significant (p-value>0.05, **Fig. 3e**), possibly due to the smaller sample size (**Fig. 1c**). Of note, along the ABA trajectory, sex- and *APOE*-dependent differences mirrored those found in the prAD trajectory, including a statistically significant but more subtle sex-*APOE4* interplay affecting tau levels (**Extended Data Fig. 4**). Together, these findings indicate that *APOE4* exerts its influence in early stages and throughout disease progression. Sex differences emerge later, primarily affecting tau but not Aβ pathology, in interplay with the *APOE* genotype.

### Accelerated accumulation of disease-associated glial cells in *APOE4* carriers

We next screened for cell subpopulations modulated by *APOE* genotype, sex, or both. We tested how the prevalence of each cell subpopulation is associated with *APOE4*, sex, and each of the three AD traits, using linear regression (confounder-adjusted; **Methods**, **Fig. 4a**). *APOE4* was significantly associated with known disease-associated cell subpopulations^23^, including the stress-responding astrocytes (Ast.10) and oligodendrocytes (Oli.7), the lipid-foamy-like microglia (Mic.13), a resilient subtype of PVALB+ inhibitory neurons (Inh.16)^23^ (FDR<0.05, **Fig. 4a**). The negatively associated subpopulations included the vulnerable somatostatin (SST) inhibitory neurons (Inh.6) and a cortical layer 2-3 excitatory pyramidal neuronal subpopulation (Exc.1) (FDR<0.05, **Fig. 4a**). Notably, these results matched the independent SHAP rankings produced by the DeepDynamics DNN (**Fig. 4a**, bottom). With respect to sex, most cell subpopulations showed no differences, but females displayed a significantly higher prevalence of the oligodendrocyte-lineage cell subpopulations (Oli.11 and OPC.1), which are enriched for mitochondrial programs^23^ (**Fig. 4a**).

**Figure 4.**
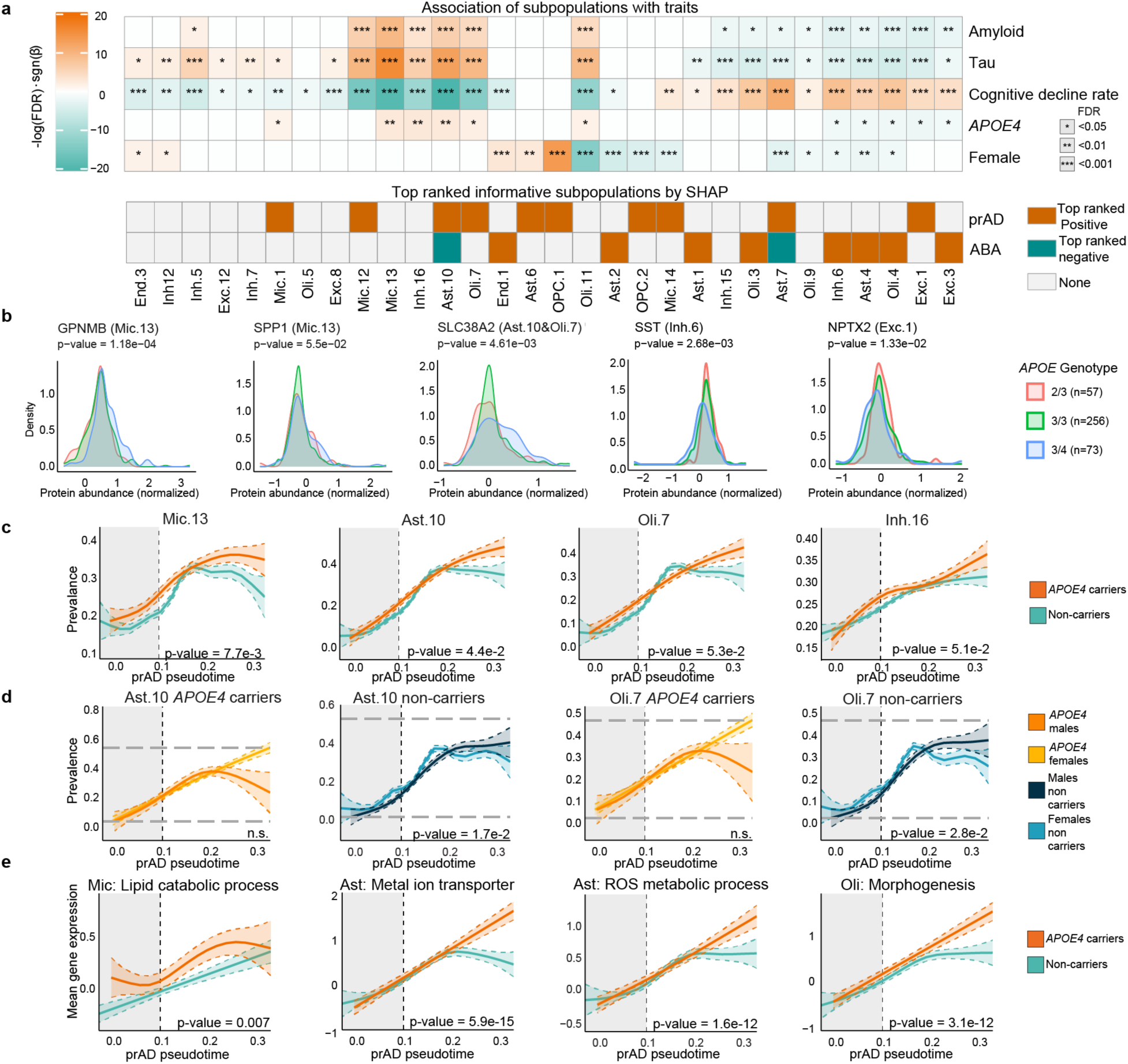
Effect of *APOE4* and sex on putative cellular drivers of AD and related molecular pathways. **(a)** Associations of cell subpopulation prevalence with AD traits (amyloid, tau, cognitive decline rate) and risk factors (*APOE4*, sex) in the bulk cohort (*n*=1,092; Linear-model: *β*=effect size; *FDR*=Benjamini-Hochberg-adjusted p-value). Bottom panel: Top-ranking subpopulations by the SHAP analysis of the DeepDynamics DNN model, colored by the directionality of the influence on the predicted prAD or ABA trajectory probability. **(b)** Validation of top-associated subpopulations by proteomics (n=400). Distribution of expression levels of known markers for each subpopulation of interest, comparing groups by *APOE genotype:* 2/3=*APOE2*/*APOE3* heterozygotes; 3/3=*APOE3* homozygotes; 3/4=*APOE4/APOE3* heterozygotes; Rare 2/4 and 4/4 samples were omitted. P-values were computed using a two-sided Wilcoxon rank-sum test. **(c-d)**, DeepDynamics-inferred dynamics of *APOE*-associated cell subpopulation prevalence along the prAD pseudotime (x-axis), stratified by: *APOE4* genotype (**c**) or combined *APOE4* and sex (**d**). (additional subpopulations in **Extended Data Fig. 5b and 6**). **(e)** Inferred *APOE4*-dependent pathway dynamics in a DeepDynamics-expanded snRNA-seq cohort (n=419). GAM-fitted dynamics of the per-individual mean pseudobulk expression of differential genes within the pathway (y-axis) along the prAD pseudotime (x-axis).

We validated the association of cell subpopulations with *APOE4* in a proteomic dataset of cortical samples from 400 individuals^41^, comparing the distributions of proteins used as markers for each subpopulation of interest, defined from their expression in the snRNA-seq atlas^23^. Indeed, the Ast.10 and Oli.7 marker protein SLC38A2, the Mic.13 markers GPNMB and SPP1, and the Exc.1 marker NPTX2 were significantly elevated in *APOE4* carriers compared to non-carriers (p-value<0.05; **Fig. 4b**). We also validated that the Inh.6 marker protein SST was significantly reduced in *APOE4* compared to *APOE3* and *APOE2*, consistent with the overall SST neuronal vulnerability in AD^23,24,29,42^. In contrast, the Mic.13 marker CPM was only marginally elevated in *APOE4* carriers (p-value<0.073), and we could not reproduce the association of *APOE4* with the Oli.7 marker QDPR and the Ast.4 marker CD44 at the protein level (**Extended Data Fig. 5a**).

We next characterized the dynamics of the 10 cell subpopulations that were significantly associated with *APOE* genotype (**Supplementary Table 3**). We found four subpopulations with significantly different dynamics in their accumulation along the prAD trajectory between *APOE4* carriers and non-carriers (p-value<0.05, **Fig. 4c; Extended Data Fig. 5b**). Specifically, disease-associated stress-responding Ast.10 (p-value<0.044) and Oli.7 (p-value<0.053) subpopulations showed accelerated accumulation in *APOE4* carriers at later stages of the prAD trajectory (**Fig. 4b**), paralleling the *APOE4*-dependent tau pathology dynamics (**Fig. 3**). A similar but more modest effect was found for the disease-associated Mic.13 (p-value<7.7e-3). Such *APOE* genotype effects were absent along the ABA trajectory (**Extended Data Fig. 6a**) and between sexes along the prAD trajectory (**Extended Data Fig. 6b**).

When examining *APOE4*-sex interplay, sex differences for the Ast.10 and Oli.7 subpopulations were weakly significant for non-carriers (0.01<p-value<0.05), possibly reflecting the smaller sample sizes in the doubly stratified subgroups (**Fig. 1c**), while female *APOE4* carriers showed a pronounced, though not statistically significant, trend of faster increase in Ast.10 and Oli.7 at later disease stages compared to male carriers (**Fig. 4d**). No strong sex-specific effect was detected in Mic.13 (**Extended Data Fig. 6c**).

Together, these results underscore the complex interplay between *APOE4*, sex, and cellular dynamics in AD progression, suggesting that specifically Ast.10, Oli.7 and Mic.13 along with specific neuronal subtypes may act as direct modulators of disease progression, especially during later pseudotimes (late disease stages).

### *APOE4*- and sex-dependent molecular signatures

Having pinpointed cell subpopulations whose prevalence is affected by *APOE4* in AD, we next investigated the *APOE4*-linked molecular changes within these cells. To obtain molecular resolution at the cell-type level, we reverted to snRNA-seq, repurposing DeepDynamics to align additional snRNA-seq samples along the prAD and ABA trajectories and thereby expanding the dataset to 412 samples (after excluding seven *APOE2/4* samples) with pseudotime annotations (**Methods, Extended Data Fig. 1f, g**).

Given the observed *APOE4*-dependent modulation of Ast.10, Oli.7, and Mic.13 dynamics, we focused on pathways enriched in the transcriptional signatures of each of these subpopulations. We modeled pathway dynamics along the prAD trajectory within their respective parent cell types: astrocytes, oligodendrocytes and microglia. For each pathway, we compiled gene sets comprising genes annotated to that pathway and present in the transcriptional signature of the relevant cell subpopulation. We then computed, per individual, the mean pseudobulk expression of each gene set. We used GAMs to model the dynamics of the pathway along the prAD trajectory within the relevant cell type (**Methods**, **Fig. 4e**). Consistent with the accelerated accumulation of these glial subpopulations (**Fig. 4b**), we observed *APOE4*-dependent upregulated expression of key pathways along disease progression (**Fig. 4e**). These pathways included accelerations of genes linked to lipid-catabolic processes in microglia (p-value<0.007), metal-ion transmembrane transporter activity (p-value<5.9e-15), the Reactive Oxygen Species (ROS) metabolic process (p-value<1.6e-12) in astrocytes, and morphogenesis in oligodendrocytes (p-value<3.1e-12, **Fig. 4e**).

We confirmed in an independent human cortical snRNA-seq SEA-AD dataset^29^ (n=82) that the signature scores for three out of the four pathways were significantly higher in *APOE4* carriers within the relevant cell types, with the exception of metal-ion transporter activity genes in astrocytes (Wilcoxon rank-sum one-sided test; p-value<0.01).

We next searched for differentially expressed genes within each *APOE4*-linked cell type Inh.6, Inh.16, Exc.1, microglia, astrocytes, oligodendrocytes cells, and potential differences between males and females (**Supplementary Table 4**). Analyzing males and females separately identified three gene categories of *APOE4* affected genes: (i) same effect in both sexes (**Fig. 5a**, top-right and bottom-left quadrants), (ii) sex-specific effect (**Fig. 5a**, along the axes), and (iii) opposing effect between sexes (**Fig. 5a**, top-left and bottom-right quadrants). To avoid biases from the over-representation of AD among *APOE4* carriers, the analysis was performed in the expanded snRNA-seq dataset separately within three subsets of individuals: healthy, prAD and ABA. In all cell types analyzed, we found an abundance of sex-specific differentially expressed genes between *APOE4* carriers and non-carriers, while microglia also harbored a large set of differentially expressed genes with the same effect in both sexes (**Fig. 5b-f**, **Extended Data Fig. 7a-e**). These results echo the sex-dependent trends observed at the cell subpopulation level for Ast.10 and Oli.7 but not Mic.13 (**Fig. 4d**, **Extended Data Fig. 6c**). Notably, *APOE4*-modulated genes were detected already in the subset of healthy individuals (**Fig. 5b-f**, **Extended Data Fig. 7a**), indicating early *APOE4*-dependent molecular modulations that might underlie the increased baseline pathology load we observed for *APOE4* carriers (**Fig. 3c**).

**Figure 5.**
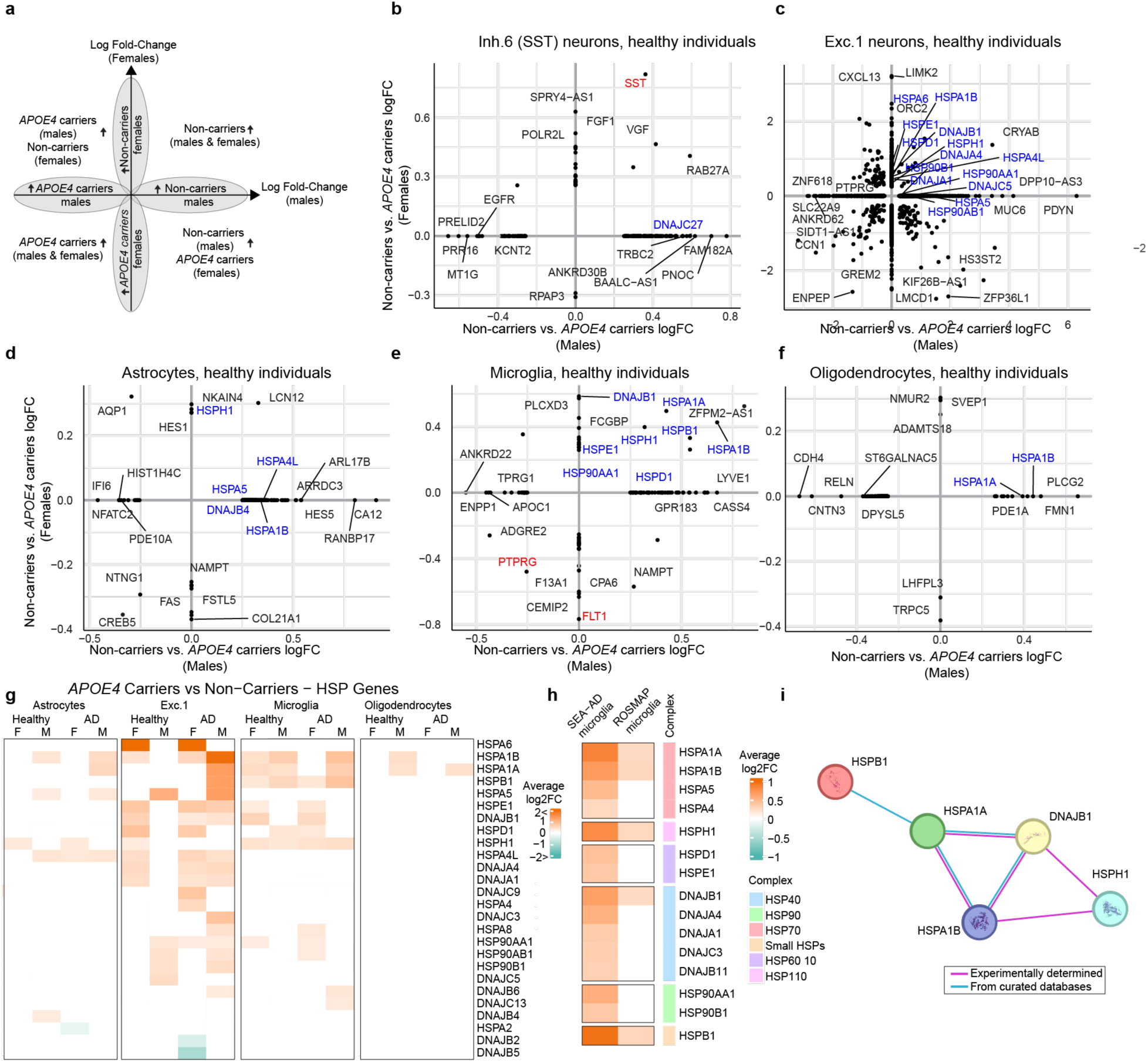
Molecular alterations in *APOE4*-associated cell types. (a-f) Sex-specific differentially expressed genes in *APOE4* carriers in healthy individuals. 2D scatter plots of differentially-expressed genes in non-carriers vs. *APOE4* carriers, in females (y-axis) and in males (x-axis) for different cell types, in healthy individuals (pseudotime < 0.1, prAD and ABA classified individuals in **Extended Data** Fig. 7). Plot interpretation in **a:** differentially expressed genes in both sexes, fall in one of the four quadrants*; sex-specific* differentially expressed genes fall on the x-axis (males) or y-axis (females). Arrows= upregulation; Blue labels mark heat shock protein (HSP) genes; red labels mark genes of interest. **(g)** Upregulated expression of HSPs in non-carriers. Heatmap of log2 fold change (non-carriers vs. *APOE4* carriers) for differentially expressed HSP genes across cell types in healthy individuals and in AD individuals. M=males, F=females. **(h)** Validation of upregulated expression of HSP genes in non-carriers compared to *APOE4* carriers in the SEA-AD independent cortical snRNA-seq dataset. Heatmap displaying the log2 fold change between non-carriers and *APOE4* carriers in expression of various heat shock protein (HSP) genes in microglia across all individuals in the SEA-AD dataset^29^ or our ROSMAP dataset. Color scale spans from -1 (turquoise, high in *APOE4*) to 1 (orange, high in non-*APOE4* carriers), no change (white). Additional cell types in **Extended Data** Fig. 8a**, Supplementary Table 5. (i)** Differentially expressed HSPs in the ROSMAP and SEA-AD microglia form an interacting network of proteins. HSP proteins (nodes) and their interactions (edges) based on STRING-DB^55^. The full HSP network in **Extended Data** Fig. 8b.

Among the differential genes, in the vulnerable Inh.6 SST inhibitory neuronal subtype, the expression of the *SST* neuropeptide gene was downregulated among *APOE4* carriers in both sexes within the healthy and ABA subsets and only in females in prAD (**Fig. 5c**, **Extended Data Fig 7b-c**). This result is consistent with our proteomic validations showing reduced SST protein in *APOE4* carriers (**Fig. 4b**). In microglia, *FLT1* was upregulated in APOE4 females in both the healthy and ABA groups, and in all APOE4 carriers in the prAD group, consistent with its role in vascular-immune signaling and amyloid-driven tau accumulation and cognitive decline^43^. Similarly, the AD-risk tyrosine phosphatase PTPRG was elevated in all APOE4 carriers (**Fig. 5e; Extended Data Fig. 7b–c**). Interestingly, across cell types in both healthy and AD individuals, we found consistent downregulation of heat shock proteins (HSPs) genes in *APOE4* carriers. Nonetheless, the specific heat-shock genes varied across cell types (**Fig. 5b-f; Extended Data Fig. 7**), and were largely sex-specific, with the highest number of upregulated genes found in male *APOE4* non-carriers (**Fig. 5g**).

We validated these findings in the independent SEA-AD cortical snRNA-seq dataset^29^ (**Supplementary Table 5**). Given the lower number of samples in this independent set we could not conduct a sex-specific analysis. Focusing on microglia cells that showed the same effect in both sexes (**Fig. 5e**), we confirmed the downregulation of multiple HSPs in *APOE4* carriers (**Fig. 5h**). We also confirmed the downregulation of HSPs in *APOE4* carriers across the other cell types excluding oligodendrocytes (**Extended Data Fig. 8a**).

Consistent with these findings, the validated HSPs within both datasets form a highly interconnected protein-protein interaction network (**Fig. 5i, Extended Data Fig. 8b**). These HSPs comprise the core HSP70 chaperones HSPA1A and HSPA1B, involved in protein aggregation clearance^44^, and their co-chaperones HSPH1, HSPB1 and DNAJB1, which stabilize and enhance HSP70 activity^45^, and facilitate proper protein folding during cellular stress^46^ (**Fig. 5i**). Of note, while expression of HSPs strongly increases along the progression of AD^21,23^, our results suggest these transcriptional changes may represent a protective cellular response, given the consistently lower expression in the high-risk *APOE4* carriers.

## Discussion

DeepDynamics bridges the high resolution of single-cell methods with the large scale of bulk datasets to chart cellular dynamics in complex biological processes, overcoming the difficulty of stratifying samples by genetic, environmental, or clinical variables. Where previous methods leveraged bulk RNA-seq chiefly to estimate cell-type composition, DeepDynamics reconstructs coupled cellular–pathological dynamics, enabling direct interrogation of disease mechanisms. Maximizing dataset size is particularly critical for linking risk factors to cellular and molecular mechanisms, an issue further amplified in temporal mapping where sparse data spread thinly across time. By jointly analyzing snRNA-seq and bulk-RNA-seq from 1,092 human brains densely sampled across the continuum of AD and aging, DeepDynamics reconstructed the cellular dynamics underlying both processes (**Fig. 1**). We identified *APOE4-* and sex-dependent modulations of cell-subpopulation dynamics and the accompanying clinicopathological states (**Fig. 3-4**). We also repurposed DeepDynamics to integrate additional snRNA-seq samples into the trajectories to provide the scale needed for exploring disease stage-specific differential expression and dynamics of molecular pathways along its progression (**Fig. 5**). While the current study centered on AD risk, DeepDynamics is a flexible framework applicable to dissecting cellular dynamics across a wide spectrum of biological systems and diseases.

Beyond resolving cell-subpopulation dynamics in bulk samples, DeepDynamics leverages explainability analysis to identify informative cellular features driving trajectory predictions (**Fig. 2**), thereby addressing another key limitation of large-scale omics studies: prioritizing putative cellular drivers of biological processes among the myriad measured cellular changes along their progression. The explainability analysis prioritized both known AD-associated subpopulations^17,21,23,29^ (*e.g.* Ast.10 and Oli.7; **Fig. 2c**) and previously overlooked, rare or subtly changing cell subpopulations (*e.g.* Peri.1, Peri.2, Oli.3, and Oli.4; **Fig.2d-f**).

DeepDynamics simultaneously tackles two challenges: measuring cell-subpopulation dynamics across disease progression and aligning those dynamics with clinicopathological traits. It scales trajectory-inference algorithms, originally devised to infer pseudotime from scRNA-seq snapshots^23,25,29^, to large-scale bulk RNA-seq cohorts, by using a DNN to assign each post-mortem brain a position along the disease timeline and reconstruct dynamics along it. Because post-mortem samples contain both molecular profiles and quantifiable pathological features, the framework aligns high resolution cellular and molecular dynamics directly with shifts in pathological and clinical traits. Other approaches to measure pathological dynamics, such as PET imaging, offer temporal resolution in living human brains but capture only coarse pathological features at a few time points, thereby yielding limited or no cell-type-specific information and providing lower quantitative precision than post-mortem measurements^47^.

Building on these capabilities, DeepDynamics offers a unified, continuous view of cell-subpopulations and clinicopathological dynamics that both recovers earlier findings and uncovers new insights into *APOE4* and sex modulation of AD risk. Specifically, DeepDynamics shows the *APOE4*- and sex-dependent acceleration of AD pathologies. Our results are consistent with previous works indicating *APOE*-sex interactions in AD risk^48,49^, yet further revealed that Aβ is modulated by *APOE4* independent of sex while tau is dependent on both. Moreover, the temporal resolution of DeepDynamics revealed that the expected accelerated accumulation of pathologies in *APOE4* carriers occurs from the earliest disease stages, followed by the expansion of specific cell subpopulations (**Fig. 3-4**). Notably, in APOE4 carriers, *the* overall cellular compositions remained unchanged, yet three disease-associated glial subpopulations, Mic.13, Ast.10 and Oli.7^23,24^ were selectively and predominantly altered (**Fig. 4c**). The perturbation of these specific glial subpopulations in *APOE4* carriers suggests they are not mere by-products of disease but active drivers of its progression, supporting prior causal computational predictions done by mediation analyses^23^. Together, these results outline a temporal hierarchy of involvement of these specific disease-associated glial subpopulations in AD progression: microglia respond first, whereas changes in astrocytes and oligodendrocytes emerge later, tightly coupled to tau accumulation in a sex-dependent manner.

Our subsequent molecular investigation links *APOE4* to distinct genes and pathways, identifying them as putative causal drivers amid the many molecular changes that accompany AD progression (**Fig. 5**). In particular, our findings indicate that the Inh.6 Somatostatin (SST) inhibitory neurons, previously identified as vulnerable in AD^23,29^, are further compromised in *APOE4* carriers, with reduced prevalence of this neuronal subpopulation and reduced abundance of the SST neuropeptide protein; SST RNA levels were already reduced in healthy *APOE4* carriers. We also uncovered a metabolic dysfunction in both microglia and astrocytes in *APOE4* carriers. Microglia of *APOE4* carriers had upregulated expression of lipid catabolic genes and the *FLT1* tyrosine kinase receptor, strengthening these pathways previously suggested as neurotoxic events exacerbating AD^43,50^. Astrocytes had higher expression of ROS metabolic genes in *APOE4* carriers consistent with previous reports in human IPS-derived *APOE4* astrocytes^51^ that we now can link to human brains (**Fig. 4**). Interestingly, across cell types APOE4 carriers show lower expression of heat-shock proteins (HSPs), consistent with reports in oligodendrocytes, including multiple HSP70 family members and their co-chaperones, which play critical roles in clearance of protein aggregations^44^ and protection against cellular stress^52,53^. Although HSPs were shown to be upregulated along the progression of AD across brain cell types^22,54^, their reduction in neurons and glial cells in APOE4 carriers suggests that impaired HSP transcription may precede, and possibly facilitate, the pathogenic cascade.

Our study introduces a framework to link risk factors to cellular and molecular dynamics in large cohorts and with single cell resolution. This approach addresses a key limitation of large-scale omics studies, which often reveal associations but not the causal drivers of disease progression. By identifying cell subpopulations, molecular pathways and specific molecules modulated by genetic risk, DeepDynamics systematically prioritizes candidate drivers of pathology that may serve as therapeutic targets, providing an analogue to perturbation experiments in model systems.

The DeepDynamics framework also has limitations. It requires overlapping snRNA-seq and bulk RNA-seq data, and a sufficient number of snRNA-seq samples to infer the initial trajectories. While this focused approached enables robust interrogation of risk factors based on validated trajectories, it may not capture other potential disease subtypes or trajectories that could be present in the larger population. Nonetheless, it offers a practical roadmap for effective experimental design: pairing a small reference cohort profiled at high-resolution with a larger cohort profiled only in bulk. Expanding the DeepDynamics framework across growing omics datasets, and informing future experiments to enhance resolution and scale, will enable rigorous mapping of risk factors to cellular functions, advance our understanding of disease mechanisms, and accelerate therapeutic target discovery.

---

**Extended Data Figure 1.**
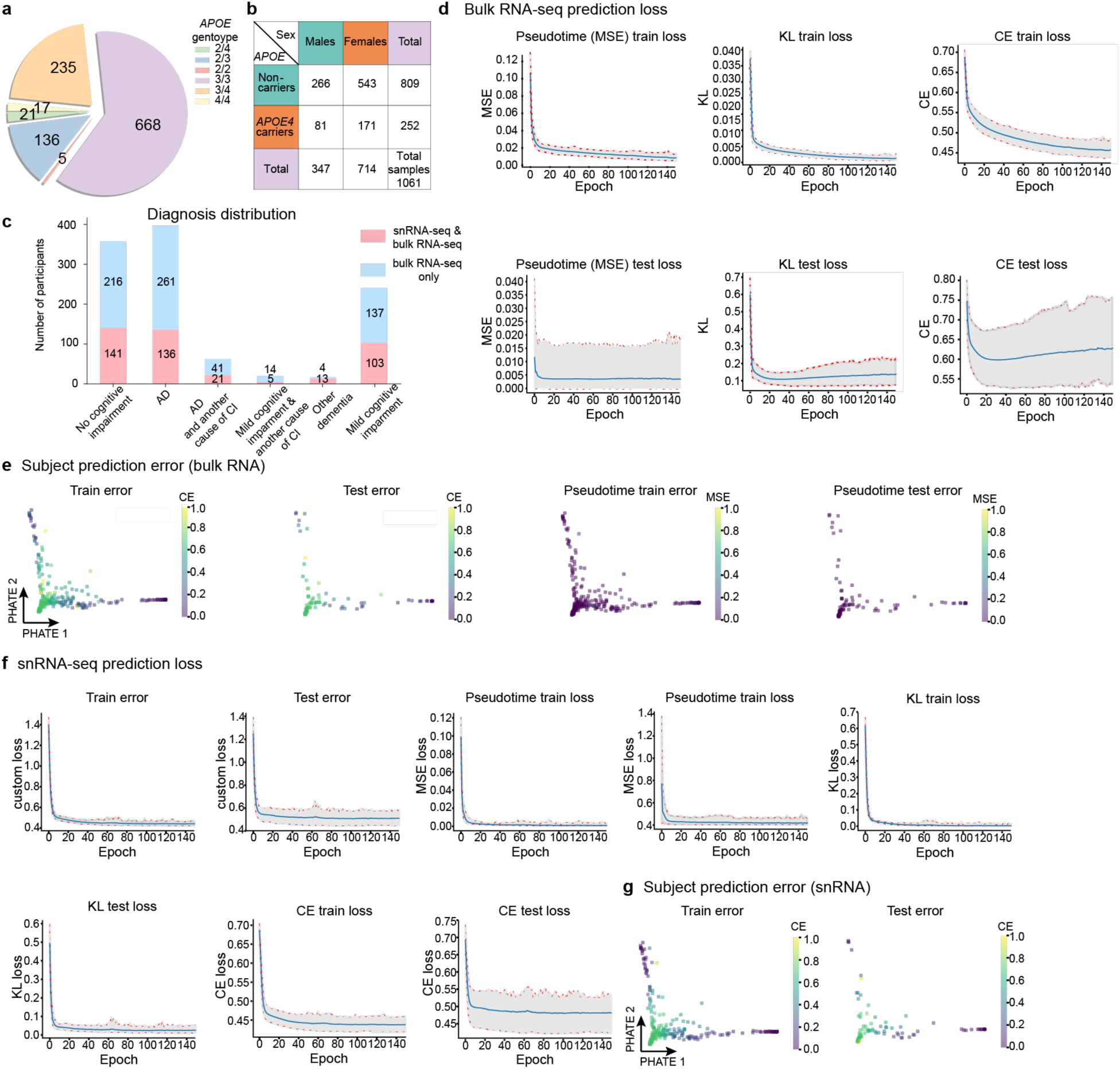
DeepDynamics and DNN model performance. **(a)** The distribution of *APOE* genotypes within our cohort. Subjects with at least one *APOE4* allele were classified as *APOE4* carriers (n=1,082). *APOE2/4* samples were excluded from the downstream analyses (n=1,061). **(b)** Distribution of samples in our study by *APOE4* carriers and sex (n=1,061). **(c)** Distribution of samples within the snRNA-seq and bulk RNA-seq (pink) cohorts by cognitive diagnosis, showing the increased coverage in the bulk RNA-seq data (blue) across categories. CI: cognitive impairment. **(d)** Training (top) and test (bottom) loss (y-axis) for the DNN model across training epochs (x-axis) for different loss measures: MSE=Mean Squared Error, KL=Kullback-Leibler divergence, CE=Cross Entropy. **(e)** Prediction error per individual. 2D embedding (PHATE^56^) of subjects for visualization, colored by: prediction error (CE loss, two right panels) or the MSE loss (two left panels) in the train and test. The CE loss decreases as the pseudotime advances in both trajectories, and the MSE loss is small across the entire landscape. **(f)** DeepDynamics can be repurposed to align snRNA-seq samples along given trajectories. Prediction and validation loss terms shown as in (d) applied on snRNA-seq, including custom loss (as in Fig. 1e), MSE, KL and CE. SnRNA-seq assignment predictions have smaller loss overall compared to bulk. **(g)** Test and train error per individual. Same as (e) for predictions from snRNA-seq data.

**Extended Data Figure 2.**
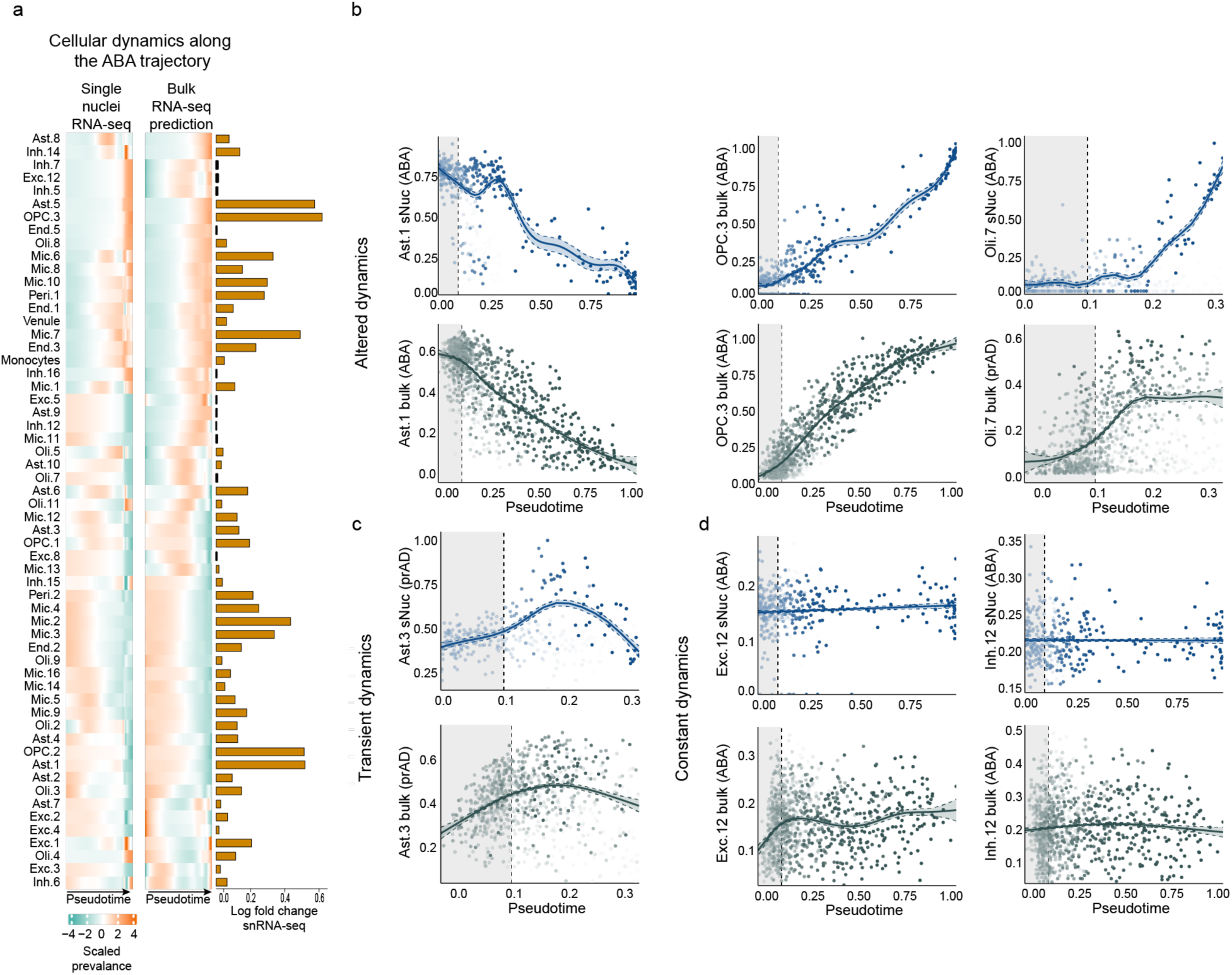
Validation of DeepDynamics predicted trajectories. **(a)** Bulk-RNA-predicted cellular dynamics match snRNA-seq. Scaled GAM estimates of prevalence (proportion) for each cell subpopulation (rows) as a function of pseudotime along the ABA trajectory (y-axis) for snRNA-seq analyzed by BEYOND^23^ and bulk RNA-seq by DeepDynamics (prAD trajectory in Fig. 2a). Horizontal bar plot (right): magnitudes of log-fold change (LFC) between minimum and maximum prevalence values for each subpopulation in the snRNA-seq dataset; Smaller LFC values indicate relatively static dynamics. **(b-d)** Representative subpopulations with different types of temporal dynamics, altered (increasing or decreasing along pseudotime, **b**), transient changes (**c**), and constant (**d**) dynamics. The GAM-fitted prevalence of each cell subpopulation (y-axis) is plotted as a function of pseudotime (x-axis) for the snRNA-seq or the bulk RNA.

**Extended Data Figure 3.**
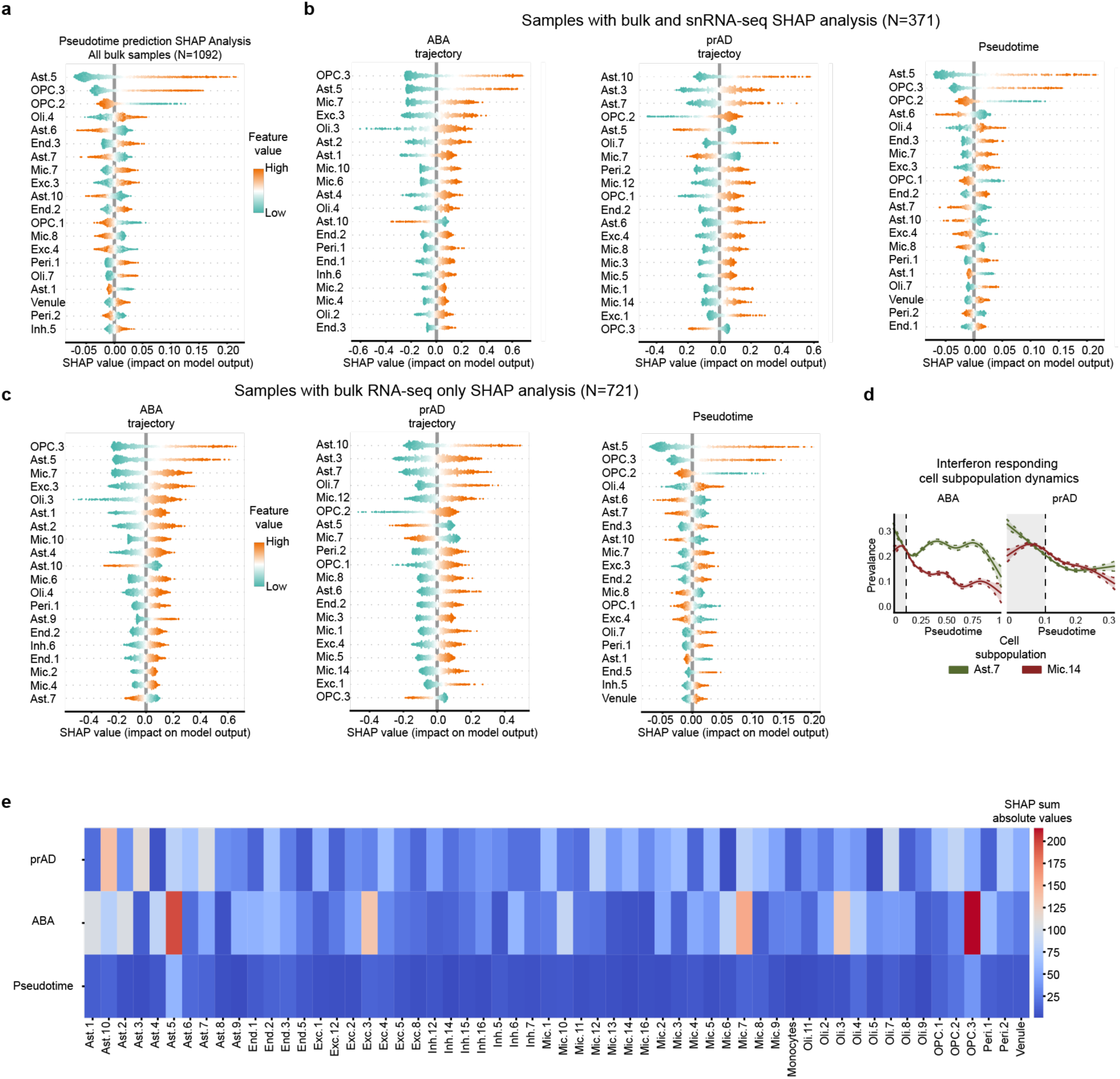
SHAP explainability analysis of the DeepDynamics DNN model. (a-c) SHAP explainability analysis identifies informative cell subpopulations. Distributions of SHAP values (x-axis) in the bulk dataset for top-ranked cell subpopulations (y-axis). Ranked by mean absolute SHAP value, quantifying the impact of each subpopulation on predictions for the: pseudotime in all bulk RNA-seq samples (**a**), trajectory probabilities and pseudotime in the bulk RNA-seq samples within the training set only (**b**), trajectory probabilities and pseudotime in the bulk RNA-seq samples within the test set only (**c**). **(d)** Dynamics of interferon-responding glial subpopulations. GAM-fitted prevalence of each cell subpopulation along the ABA (left) and prAD (right) pseudotime. Dashed lines = confidence intervals. **(e)** SHAP sum absolute values over all samples per cell subpopulation.

**Extended Data Figure 4.**
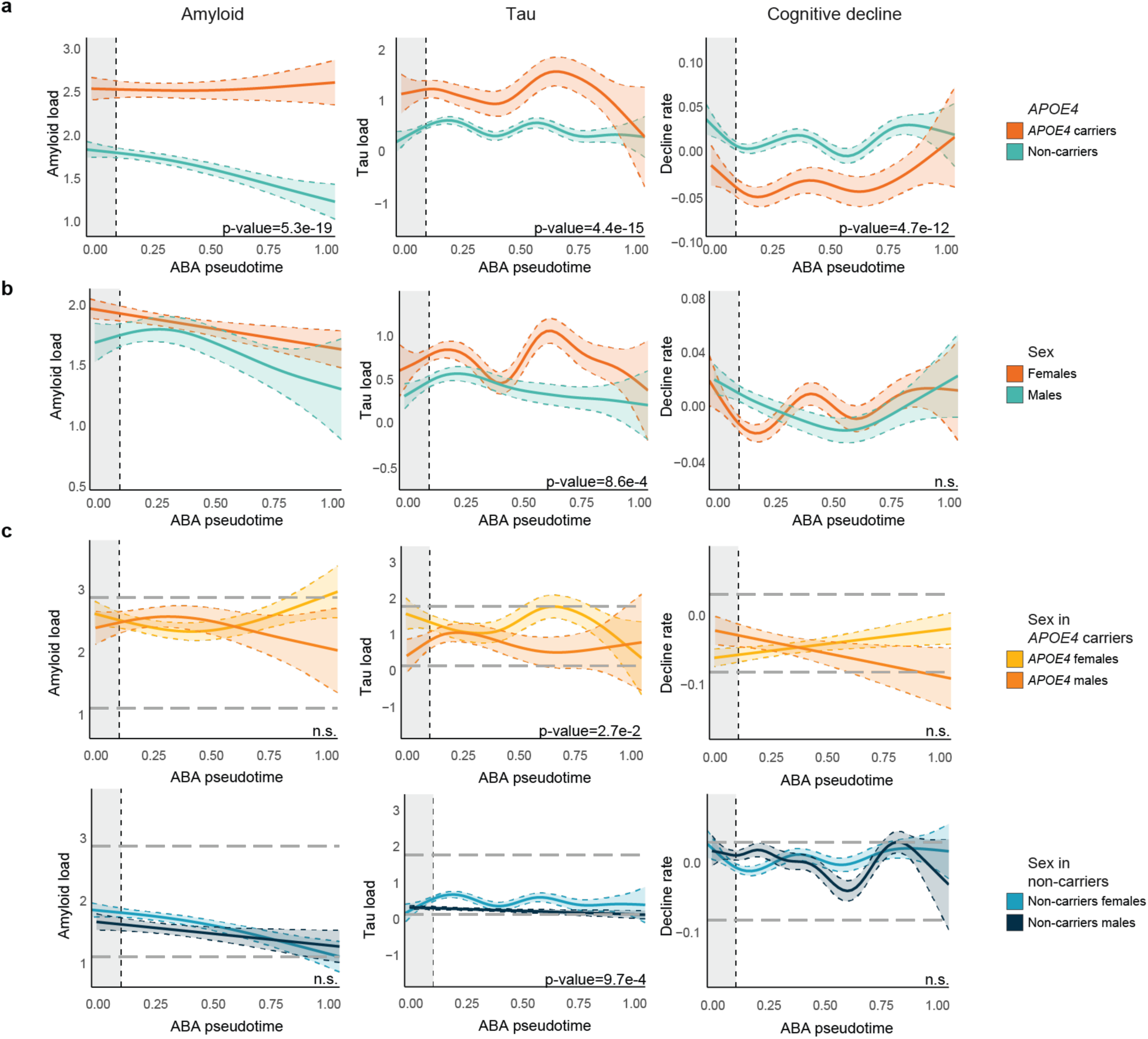
*APOE4* and sex-dependent modulation of dynamics of AD clinicopathologies. (**a-c**) DeepDynamics-inferred AD clinicopathological dynamics along the ABA pseudotime (x-axis, n=1,092, prAD trajectories shown in Fig. 3c**-e**), stratified by: *APOE4* genotype (**a**), sex (**b**), or combined stratification by *APOE4* and sex (**c**). n.s. = non-significant.

**Extended Data Figure 5.**
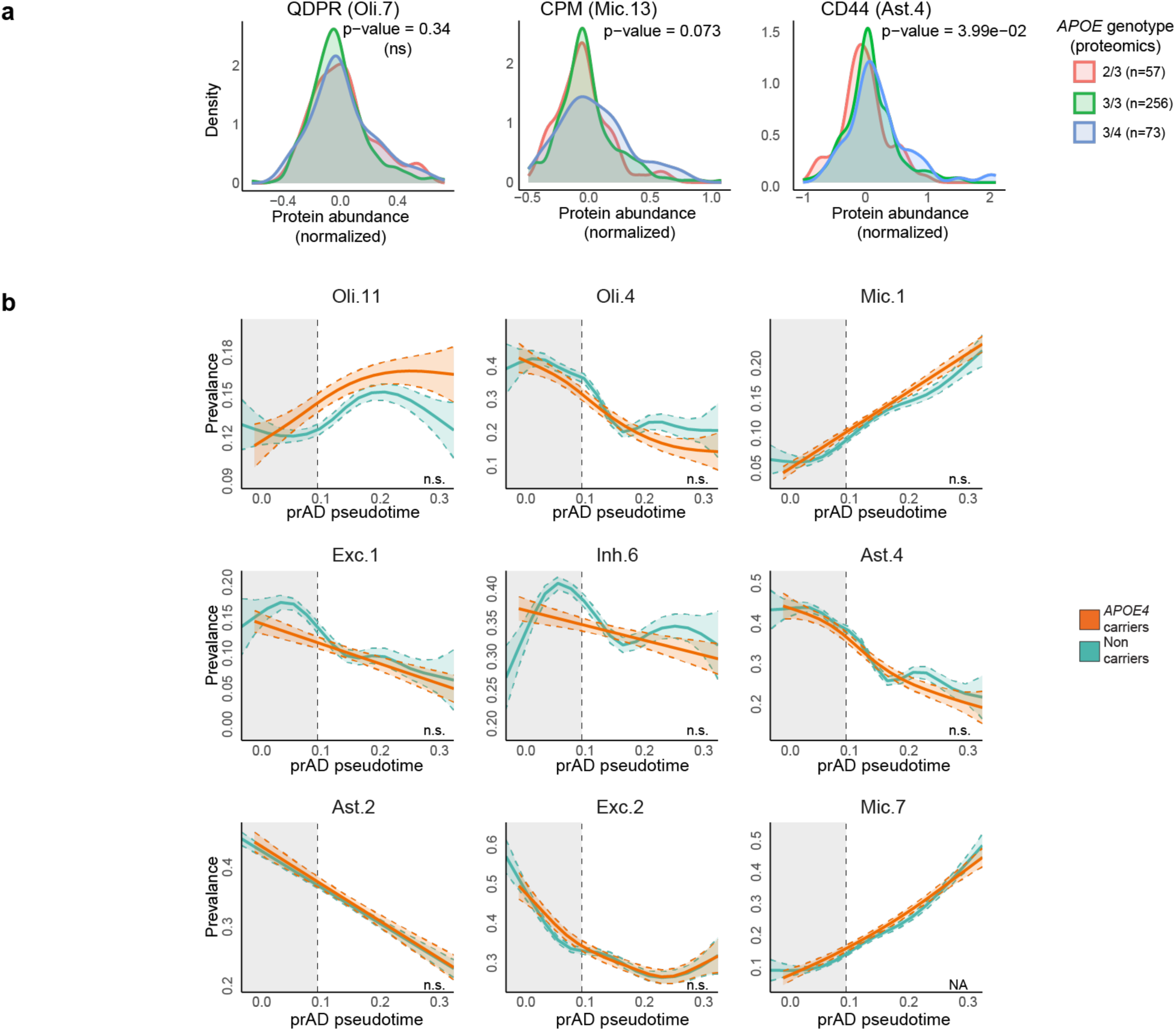
Effect of *APOE4* on cell subpopulations and dynamics along AD progressions: **(a)** Validation of top-associated subpopulations by proteomics (n=400). Distribution of expression levels of known markers for each subpopulation of interest, comparing groups by *APOE genotype:* 2/3=*APOE2*/*APOE3* heterozygotes; 3/3=*APOE3* homozygotes; 3/4=*APOE4/APOE3* heterozygotes; P-values were computed using two-sided Wilcoxon rank-sum test. Including CPM as Mic.13 marker, QDPR as Oli.7 marker, and CD44 Ast.4 marker. **(b)** Inferred dynamics of *APOE*-associated cell subpopulation prevalence (y-axis) along the prAD pseudotime (x-axis) in the bulk RNA (n=1,092), stratified by *APOE4* genotype (accompanying subpopulations in Fig. 4c).

**Extended Data Figure 6.**
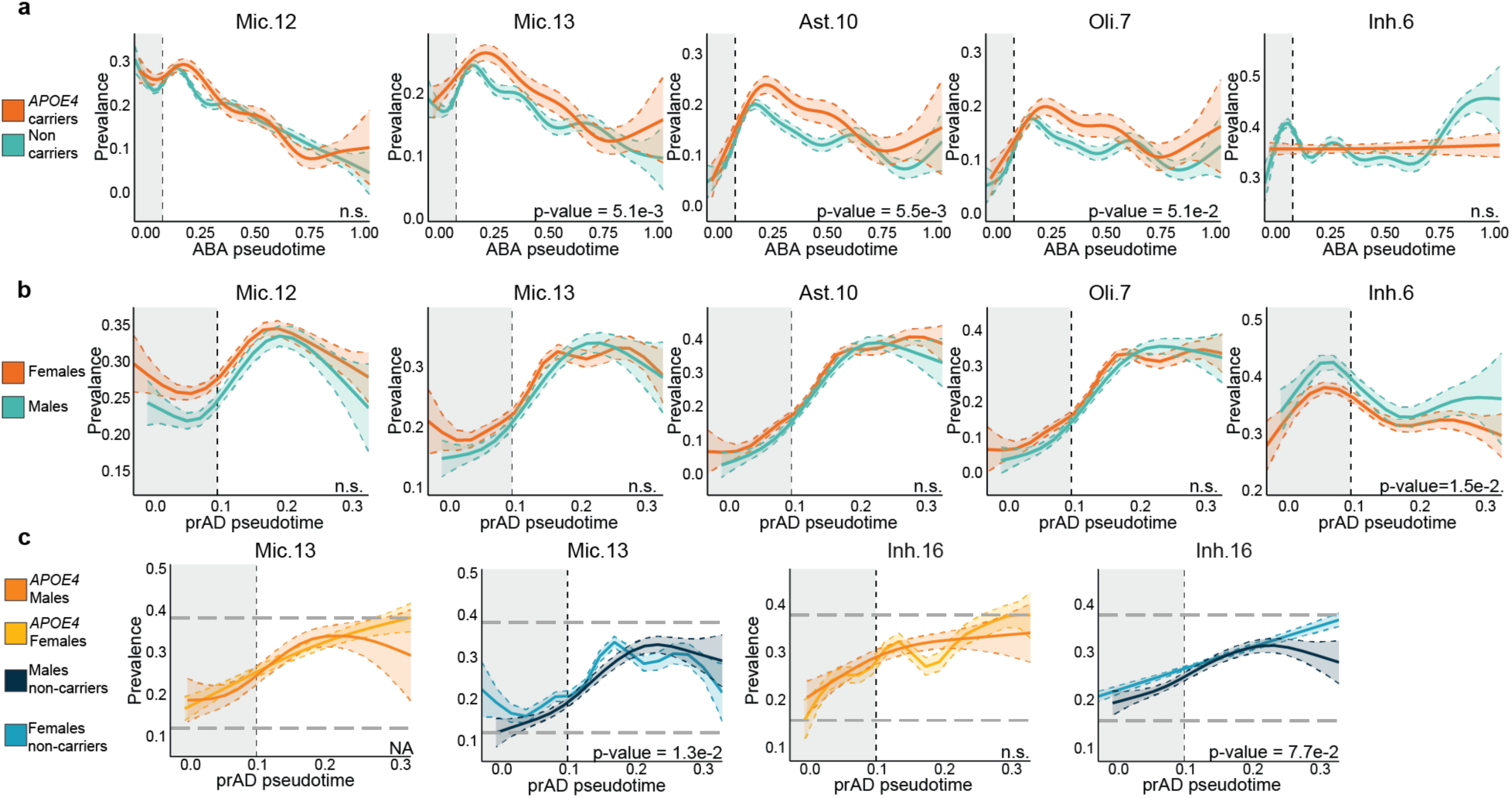
*APOE4*-modulated dynamics of cell subpopulations and molecular pathways. (**a-c**), Inferred dynamics of *APOE*-associated cell subpopulation prevalence (y-axis) along pseudotime (x-axis), stratified by: APOE4 genotype along the ABA trajectory (**a**), sex along the prAD trajectory (**b**), or combined *APOE4* and sex along the prAD trajectory (**c**). (accompanying Fig. 4c**-d**).

**Extended Data Figure 7.**
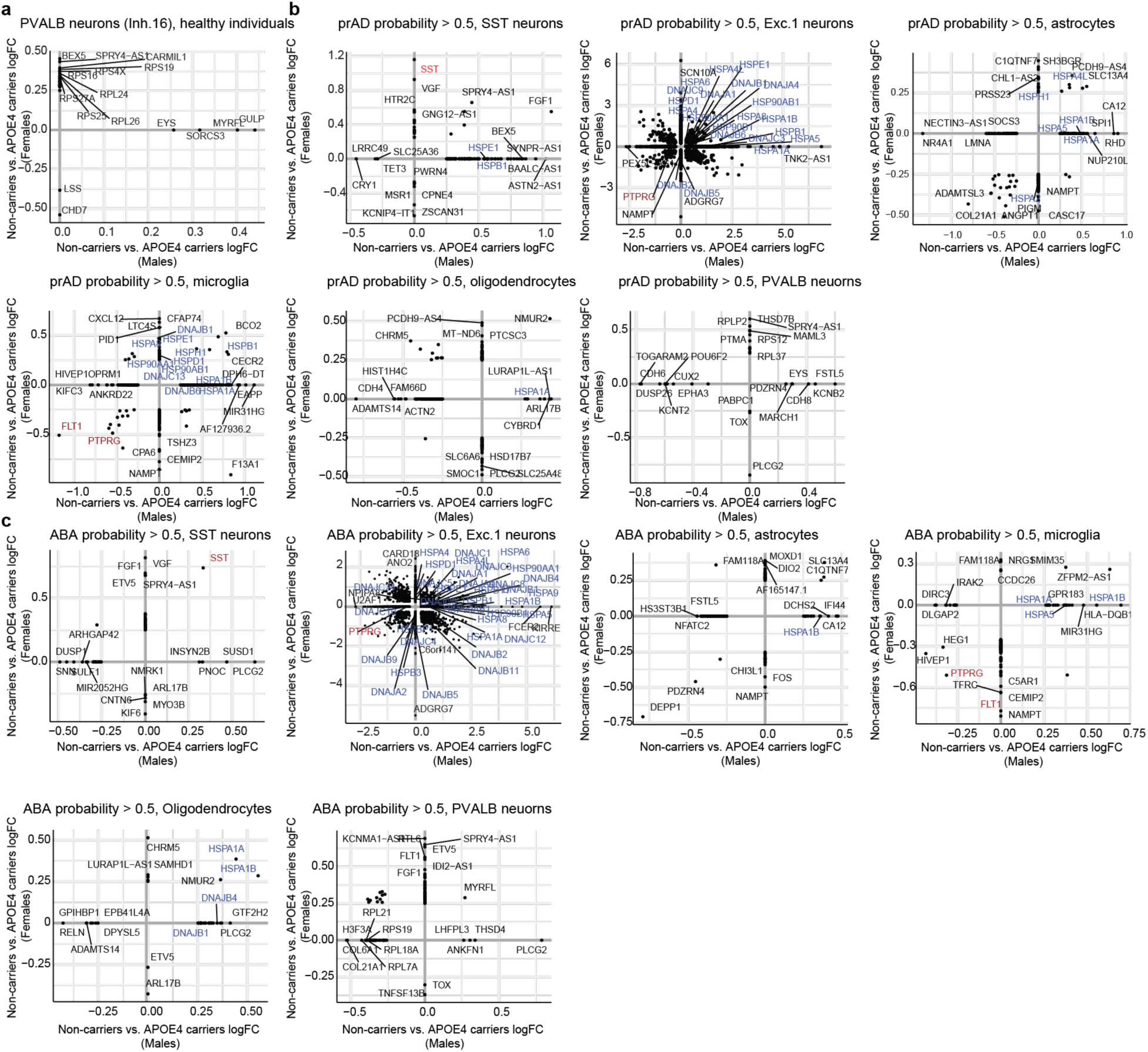
Differentially expressed genes in *APOE4* carriers within subsets of aging individuals. (**a**) Sex-specific differentially expressed genes in Inh.16 PVALB inhibitory neurons in *APOE4* carriers. 2D scatter plots of differentially-expressed genes in non-carriers vs. *APOE4* carriers, in females (y-axis) and in males (x-axis) for Inh.16 PVALB inhibitory neuronal subtype, in healthy individuals (pseudotime < 0.1). **(b-c)** Sex-specific differentially expressed genes in *APOE4* in individuals classified within the prAD (**b**) and ABA (**c**). Presented as in a. prAD individuals defined as pseudotime > 0.1 and prAD probability >0.5; ABA individuals defined as pseudotime > 0.1 and ABA probability > 0.5.

**Extended Data Figure 8.**
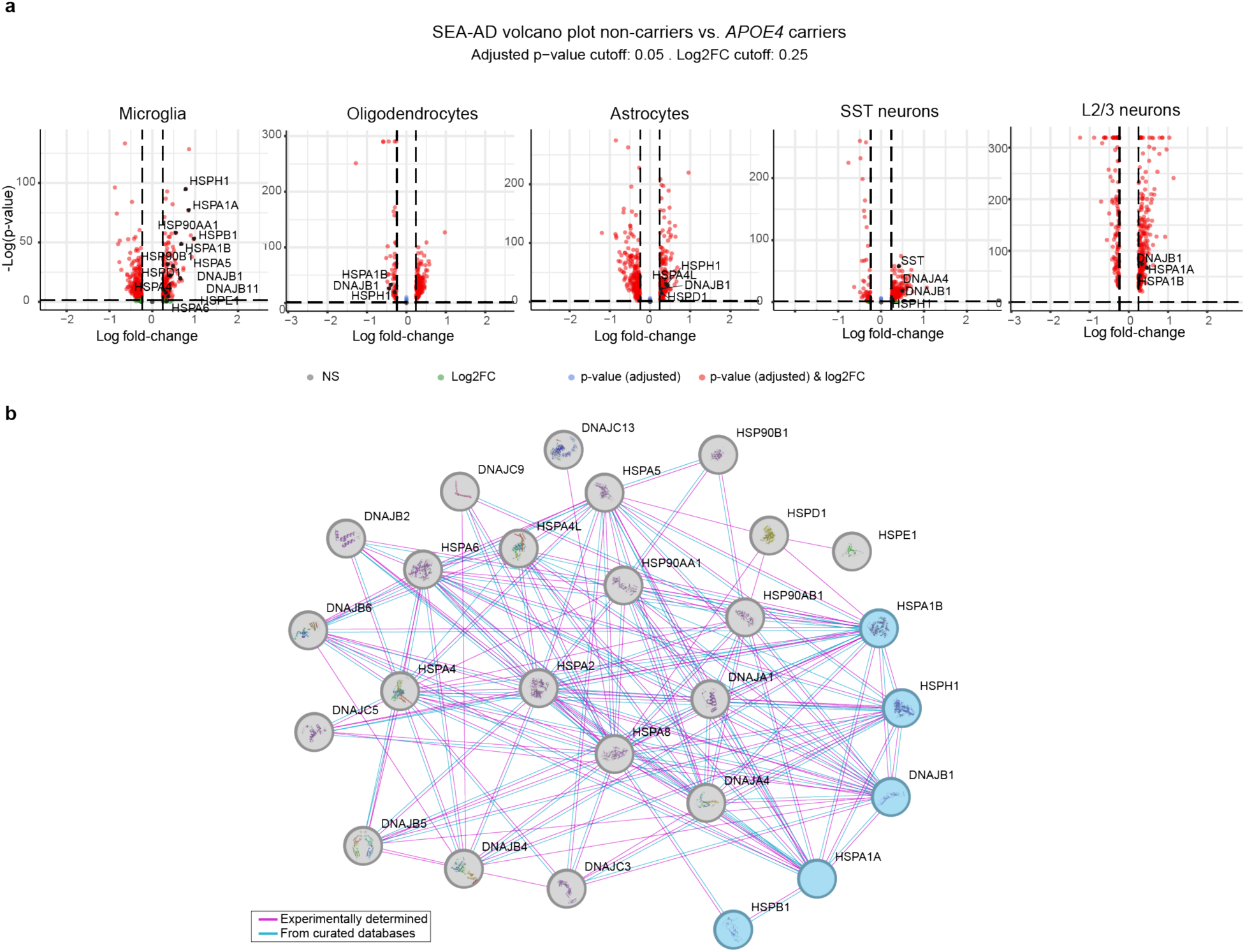
Validation of differential gene expression in *APOE4* in an independent Seattle-AD dataset: **(a)** Validation of HSP differential expression in APOE4 across cell types. Volcano plot showing differentially expressed genes between *APOE4* non-carriers vs. *APOE4* carriers in five *APOE4*-linked cell types (microglia, oligodendrocytes, astrocytes, SST neurons, L2/3 neurons) from the independent SEA-AD snRNA-seq human cortical dataset. The x-axis represents the Log2 fold change, and the y-axis represents the -Log10 adjusted p-value. The adjusted p-value cutoff is 0.05, and the Log2FC cutoff is 0.25. Positive log fold change is upregulated in *APOE4* non-carriers. Differentially expressed HSPs form an interacting network of proteins. Protein-interactions (from STRING-DB^55^) of differentially expressed HSP proteins found in the ROSMAP cohort across cell types. Blue color = HSPs upregulated in *APOE4* non-carriers both in the ROSMAP and the SEA-AD datasets in microglia cells.

## Methods

### Experimental design - ROSMAP dataset

We applied the DeepDynamics pipeline to data obtained from participants in the Religious Orders Study (ROS)^35^ and the Rush Memory and Aging Project (MAP)^34^, two longitudinal clinical-pathologic cohort studies of aging and dementia, collectively referred to as ROSMAP. All the participants are without known dementia at enrollment, underwent annual clinical evaluations, and agreed in advance to brain donation at death. At death, the brains undergo a quantitative neuropathologic assessment, and the participant’s rate of cognitive decline is calculated from the longitudinal cognitive measures that include up to 25 yearly evaluations^63^. Each study was approved by the Institutional Review Board of Rush University Medical Center. All the participants signed an informed consent, Anatomic Gift Act documentation, and repository consent. The scRNA-seq dataset comprises 437 participants, blinded to their neuropathologic and clinical traits, and based on availability of frozen pathologic material from the dorsolateral prefrontal cortex (DLPFC, BA9), including only participants with RNA integrity number (RIN) > 5 and post-mortem interval (PMI) < 24 h, as in our previous studies^23,25^, yielding 1.68 million single-nucleus RNA profiles from neuronal and glial cells, fully annotated to cell types and cell subpopulations in Green *et al*^23^. Out of these 437 participants, 386 participants were aligned to two aging trajectories: progression of AD (prAD) and Alternative Brain Aging (ABA). The bulk RNA-seq dataset comprises 1,092 individuals (419 overlapping with snRNA-seq). The bulk proteomics dataset comprises 400 individuals (146 overlapping snRNA-seq, 400 overlapping bulk RNA-seq), which we used for validation. Proteomic characterization utilized mass spectrometry, quantifying > 8,600 proteins^41^.

Clinicopathological measures were collected as part of the ROSMAP cohorts (previously described^57–59)^. We focused our analysis on three quantitative AD traits: amyloid-β (Aβ) load, tau load, and cognitive decline rate.

#### Aβ and tau loads

The quantification and estimation of the burden of parenchymal deposition of Aβ and the density of abnormally phosphorylated tau-positive neurofibrillary tau levels present at death (which we refer to as Aβ and tau pathology, respectively) were done as follows. Tissue was dissected from eight regions of the brain: the hippocampus, entorhinal cortex, anterior cingulate cortex, midfrontal cortex, superior frontal cortex, inferior temporal cortex, angular gyrus and calcarine cortex. Sections (20 µm) from each region were stained with antibodies against the Aβ and tau proteins, and quantified using image analysis and stereology. Measurements were summarized to provide a global measure of Aβ and tau burdens. For the trait association and dynamics analyses, we used the measurements of Aβ and tau in the midfrontal cortex. The load of these pathologies at the midfrontal cortex is a potentially better proxy for the pathology in the DLPFC brain region compared with the pathology load across the entire brain. Furthermore, we note that we are assessing cellular changes in the fresh-frozen DLPFC samples from one hemisphere of each brain relative to the measures of cortical Aβ and tau load measured in the midfrontal cortex of the opposite, fixed hemisphere where the standard, structured neuropathologic assessment is conducted.

#### Rate of cognitive decline

This trait was quantified using uniform structured clinical evaluations, including a comprehensive cognitive assessment administered annually to the ROS and MAP participants. The cognitive assessment included 19 cognitive performance tests, 17 of which were used to obtain a summary measure for global cognition as well as measures for five cognitive domains of episodic memory, visuospatial ability, perceptual speed, semantic memory and working memory. The summary measure for global cognition is calculated by averaging the standardized scores of the 17 tests, and the summary measure for each domain is calculated similarly by averaging the standardized scores of the tests specific to that domain. The ROS and MAP methods of assessing cognition have been extensively summarized in previous publications^60–62^ . Further statistical details have been previously described^63^.

### The DeepDynamics pipeline

#### Preprocessing of dataset

The preprocessing stage for the DNN input and labels proceeds as follows: The model input includes the prevalence of cell subpopulations inferred by RNA signatures within the tissue of interest (in our case bulk RNA-seq). These can be calculated directly when using sc/snRNA-seq data, or inferred for bulk RNA-seq data, using clustering analysis or co-expression gene module analysis in an overlapping sc/snRNA-seq atlas (e.g. CelMod^24^, see below). The model labels are the assignments of samples to trajectory probabilities and pseudotime using algorithms such as BEYOND^19^.

### Cell subpopulation deconvolution inference from Bulk RNA-sequencing

We used CelMod^24^ to determine the cellular composition of each individual bulk RNA sample. CelMod relies on a consensus of gene-wise regression models, with cross-validation to estimate accuracy. We chose the threshold for CelMod p-value to be 0.005, in order to only include the most robust cellular states, while not limiting the DNN model information (**Supplementary Table 1)**.

### DNN architecture and training

The input, output, and training labels for the DNN are as follows:

*Input:* Predicted cell subpopulation prevalence for each bulk sample (inferred here using CelMod, but can be substituted with similar algorithms^64,65^)

*Output:* Probabilities for the progression of AD (prAD) and Alternative Brain Aging (ABA) trajectories, and predicted pseudotime per sample.

Training labels: Trajectory probabilities and pseudotimes for the training samples calculated from snRNA-seq measurements (here using BEYOND, but can be substituted with similar trajectory-inference algorithms).

Unlike typical probabilities, the trajectory probabilities predicted by the model are assigned independently to each trajectory, meaning that they are not required to add up to one. As a result, samples may potentially belong to both or neither trajectory. This design is less affected by the sample size and potentially allows the model to capture more complex biological states, specifically enabling the flexibility to capture multiple parallel biological processes, such as alternative paths of aging and disease.

The DNN itself consists of linear layers with ReLU activation functions. We experimented with a range of hyperparameters, as follows:

- DNN depths: 3 - 7 layers
- Learning rate: 0.01 - 0.0001
- Optimizer: Adam
- Batch size: 5 - 20
- Early stopping: Patience of 10 epochs
- 𝛿 values: e^-6^ - e^-3^

Here, the final default hyperparameters we chose were 4 layers, learning rate of 1e^-5^, batch size of 10, and 𝛿 value of 5.5e^-5^.

The model loss function incorporated three metrics:

1. Cross Entropy (CE): Local loss for sample probability prediction.
2. Kullback-Leibler (KL) divergence: Global loss to maintain dataset-wide probability distribution properties.
3. Mean Squared Error (MSE): Loss for pseudotime prediction.

More formally:

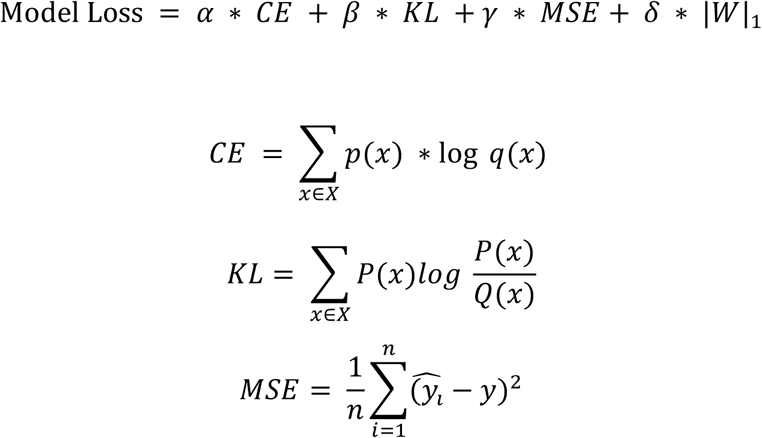

Where α, β, and γ control model balance between local probability prediction, global probability prediction, and pseudotime prediction, and δ is an L1-regularization term applied to model weights (*W*).

Training data was split 75:25 for train and test datasets, respectively. Model hyperparameters were optimized using 5-fold cross-validation. Bayesian optimization was employed to efficiently search the hyperparameter space using the *Ray Tune* package^66^.

### SHAP analysis

We used Kernel SHAP analysis to explain the model outputs: trajectories probabilities and predicted pseudotime. We tested the results for different subsets of the data: only sNuc + bulk subjects, bulk only subjects, and all the bulk data, deeming the results as robust to these compositional changes (**Extended Data** Fig. 3). The SHAP analysis and plots were done using the *SHAP* package (version 0.47.1).

### Statistical analysis of trait association

Association analysis was done using the *lm* function from *stats* R package (version 4.3.1), which is used to fit linear models with the following formula: *trait ∼ covariate + control*. The direction of the association and its significance were determined by the beta and p-value, which are the outputs of the fitted linear model. P-values were then corrected for multiple hypothesis testing (*p.adjust* from *stats* package, method = “BH”).

#### Association of AD-traits to genetic risk factors

Statistical associations between AD-related traits and the *APOE* genotype and sex risk factors were tested by regression of the traits on the binary annotation of the risk factor of interest. To remove confounding effects, we adjusted for age of death. To avoid confusion, we excluded *APOE2/4* carriers from all comparative analyses.

#### Association of AD-traits to cell subpopulations

When associating the prevalence of cell subpopulations, in addition to age of death we also control for RNA Integrity Number (RIN) and Post-Mortem Interval (PMI).

### Fitting Generalized Additive Models to trait dynamics along trajectory pseudotimes

To capture the dynamics of specific cell subpopulations and associated clinicopathologies along the defined trajectories, we fitted General Additive Models (GAM) using the R package *mgcv*, which is specifically designed for the computation of mixed GAMs with automatic smoothness estimation.

In our analysis, we utilized spline fitting to construct a generalized additive model for a given feature value *y* along a trajectory *j*:

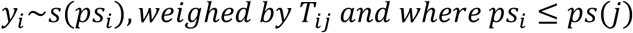

Where *i* denotes an individual, 𝑦_𝑖_ is the feature 𝑇_𝑖\_ is the trajectory probability for individual *i* and trajectory *j*. The final dynamics were predicted over equidistant pseudotime values in the range [0, ps(j)] (*mgcv::predict.gam, se.fit=TRUE*). When visualizing these dynamics, we displayed the predicted feature values across the equidistant pseudotime intervals, accompanied by a confidence interval area representing predicted values ±2 times the standard error (se), as derived from the *predict.gam* output. The computed dynamics for the pathologies and cell subpopulations are provided in **Supplementary Table 2**.

### Dynamics of cell-type-specific molecular pathways

To investigate the dynamics of key molecular pathways, we focused on pathways previously identified as enriched and specifically upregulated in disease-associated subpopulations of interest compared to all other subpopulations of the same cell type, as found in the snRNA-seq dataset^23^. To infer the pathway dynamics, we first calculate the pathway expression score (signature) for the pathway within the relevant cell type for each individual as follows:

For each individual *i*, and gene *g* within the differentially expressed genes of the pathway of interest, we compute the average expression of the gene (denoted as 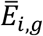) across all cells from this cell type within this individual (pseudobulk expression):

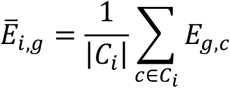

where 𝐶_𝑖_ is the set of cells for individual 𝑖.

We then standardized gene expression using a z-score transformation:

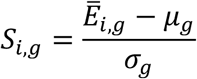

where 𝜇_b_ and 𝜎_b_ represent the mean and standard deviation of gene 𝑔 across all individuals, respectively.

Once we obtain the scaled expression values 𝑆_𝑖,*g*_, we next derived a pathway expression score for each individual *i*, (denoted as 𝑃_𝑖_) by averaging the scaled expression of all genes within a given enriched pathway 𝑃:

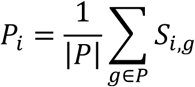

Finally, we fit the dynamics of each pathway expression over pseudotime using the same GAM-based framework described above. The computed dynamics for pathway score are provided in **Supplementary Table 2**.

### ANOVA test of Risk Group Stratification

To assess the contribution of risk group stratification to modeling disease dynamics for AD-traits, cell subpopulation prevalence or expression of a gene-set within a known pathway, we employed an ANalysis Of VAriance (ANOVA) framework based on a likelihood ratio test. Specifically, we compared a null GAM model, which did not include risk stratification (i.e. 𝑦 ∼𝑠(𝑝𝑠𝑒𝑢𝑑𝑜𝑡𝑖𝑚𝑒)), to an interaction model that incorporated risk group effects (i.e. 𝑦∼𝑟𝑖𝑠𝑘 + 𝑠(𝑝𝑠𝑒𝑢𝑑𝑜𝑡𝑖𝑚𝑒, 𝑏𝑦 = 𝑟𝑖𝑠𝑘)). This comparison was performed using the *anova* function in R with the argument *test = “Chisq”*, which yields a likelihood-ratio *χ²* statistic on the difference in model deviances. A statistically significant improvement in model fit (defined as p-value < 0.05), indicated that the inclusion of risk group stratification provided additional explanatory power. In such cases, we considered the stratification by risk groups to offer meaningful and informative value for understanding disease dynamics. The ANOVA p-values corresponding to the dynamics of the various risk presented in this study are provided in **Supplementary Table 3**.

### Differential Gene Expression Analysis

Within a given cell type, differentially expressed genes (DEGs) were computed between the low-risk control group to the high-risk group (*APOE4* carriers, excluding *APOE2/4* samples) using a *Poisson* test with a designated function from Seurat R package v5.0, using a custom mean function for the calculation of the fold change:

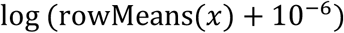

Only genes that are expressed in at least 10% of the cells with at least 0.25 log fold-change were considered.

We compared *APOE4* carriers to non-carriers separately in males and females, and within the following four groups of individuals: (1) all participants within the cohort, (2) healthy groups (pseudotime < 0.1), (3) AD (defined as individuals with pseudotime > 0.1 and prAD probability > 0.5), and (4) ABA (defined as individuals with pseudotime > 0.1 and ABA probability > 0.5). The results were corrected for multiple hypothesis using the FDR Benjamini-Hochberg correction, with a threshold of 5% FDR. The analysis results for each group and cell type presented in this study are provided in **Supplementary Table 4**.

### SEA-AD snRNA-seq dataset analysis

We validated the downregulated expression of heat shock proteins in *APOE4* carriers, and the upregulated expression of selected pathways in *APOE4* carriers in an independent snRNA-seq dataset from the SEA-AD aging brain cohort in which the middle temporal gyrus (MTG) brain region was profiled^29^.

For the differential expression analysis, we selected cells of the *APOE4*-associated cell types: oligodendrocytes, microglia, astrocytes, L2/3 neurons and SST neurons, and performed differential gene expression analysis between *APOE4* carriers (3/4 and 4/4 genotypes) and non-carriers (2/2, 2/3 and 3/3 genotypes). We used the MAST test implemented in *Seurat*. Consistent with our previous criteria, only genes that are expressed in at least 10% of the cells and at least 0.25 log fold-change are considered. The data processing of the dataset: the h5ad file from SEA-AD cohort^29^ was split by cell types, and *Seurat* objects were generated using *SeuratObject* package, using the normalized counts and cell metadata. Due to *Seurat*’s object size limitations (with the L2/3 neurons object comprising approximately 330,000 cells and 36,000 genes, exceeding the 2^31 sparse matrix size limit), we divided the L2/3 neuron data into six smaller objects, each containing up to 6,505 genes. Differential expression analysis was conducted on each subset, and the results were subsequently combined, with multiple hypothesis testing correction applied across all identified genes (*p.adjust, method = “fdr”*).

To validate the cell-type-specific *APOE4*-differential pathways, we calculated the pathways score in SEA-AD cohort within the relevant cell type using the same approach as in our main cohort. We then performed a one-sided Wilcoxon rank-sum test to assess whether pathway scores were higher in the *APOE4* carriers (high risk) group compared to the non-carriers group. The results of these analyses for each cell type in the SEA-AD cohort are provided in **Supplementary Table 5**.

## Data availability

Datasets used in this study are available at the AD Knowledge Portal (https://adknowledgeportal.org). The AD Knowledge Portal is a platform for accessing data, analyses and tools generated by the Accelerating Medicines Partnership (AMP-AD) Target Discovery Program and other National Institute on Aging (NIA)-supported programs to enable open-science practices and accelerate translational learning. Data are available for general research use according to the following requirements for data access and data attribution (https://adknowledgeportal.org/DataAccess/Instructions). Access to the content described in this Article is available online:

Synapse database for raw and processed snRNA-seq data (https://www.synapse.org/#!Synapse:syn31512863);

Synapse database for bulk RNA-seq dataset (https://www.synapse.org/#!Synapse:syn3388564); Other ROSMAP resources can be requested at the RADC Resource Sharing Hub (https://www.radc.rush.edu).

## Code availability

The complete code base used in this study is available at GitHub (https://github.com/naomihabiblab/DeepDynamics) and includes the full DeepDynamics pipeline, including a tutorial python notebook demonstrating how to move from an annotated snRNA-seq cohort (step 1a) with trajectories (step 1b), and a matching deconvolved bulk RNA-seq cohort (step 2) to bulk trajectory prediction (step 3) and cell subpopulation prioritization (step 4).

## Acknowledgements

We thank the individuals who donated their brains to research through the Rush University Alzheimer’s Disease Center. This work was supported by the Israel Science Foundation (ISF) research grant no. 1709/19, the European Research Council grant 853409, the Chan Zuckerberg Initiative (CZI) Collaborative Pairs grant CP2-1-0000000058 and the Myers Foundation (N.H.); The Minerva center grant on Cell Intelligence, Israel Science Foundation grant 385/24 (B.R.); and HUJI Data Science Center (CIDR) Seed Grant (B.R., N.H). Y.R. is supported by the Beker scholarship for excellence graduate students in cognition and brain science.

## Author contributions

N.H., B.R., Y.H. and Y.R. conceived and designed the study. Y.H. and Y.R. performed the computational and statistical analyses with help from A.C.; Y.R. developed the DeepDynamics algorithms with guidance from G.G.; N.H. and B.R. provided guidance, data interpretation and supervised the work. N.H., B.R., Y.H. and Y.R. wrote the manuscript with critical comments from the co-authors.

## Supplementary Tables

**Supplementary Table 1. Cohort description:** Clinicopathological characteristics, APOE genotype and sex across participants. Inferred cell-subpopulation prevalence for each RNA-seq sample (CelMod algorithm).

**Supplementary Table 2. DeepDynamics DNN predictions:** Predictions of DNN values per participant from bulk RNA-seq. The GAM inferred dynamics for AD-associated traits and cell subpopulations along the pseudotime for subgroups stratified by: APOE genotype, sex, or APOE×sex. Dynamics of cell-type-specific pathways stratified by APOE genotype in snRNA-seq.

**Supplementary Table 3. Statistical analysis of dynamics:** ANOVA results comparing bulk RNA-seq dynamics of AD associated traits and cell subpopulation prevalence across pseudotime stratified by APOE genotype, sex, or APOE×sex, or pathway dynamics stratified by APOE4.

**Supplementary Table 4. Differential expression analysis.** Sex-specific differential-expression analysis per cell type of interest comparing APOE4 carriers vs. non-carriers in four subgroups of individuals: healthy, prAD, ABA, or all individuals within the cohort. **Supplementary Table 5. Validations in the SEA-AD cohort**: Differential expression analysis comparing APOE4 carriers vs. non-carriers per cell type of interest and statistical analysis of differential expression of pathway scores.

