## Supplementary Table 5 for "DeepDynamics resolves cell-subtype and clinicopathological dynamics from bulk RNA-seq to identify mediators of Alzheimer’s disease risk"

**This file contains supplen**

| Sheet |
|---|
| APOE4 - 2 groups |
| Pathways score |

**nentary tables as described below.**

| Description |
| --- |
| Seattle AD dataset differential expression genes analysis between the APOE group carreris vs. no-carriers |
| Validation of pathways - Wilcoxon rank sum one-sided test p-values for chosen gene sets, Hypothesis - risk is higher than no risk |
