## Supplementary Table 4 for "DeepDynamics resolves cell-subtype and clinicopathological dynamics from bulk RNA-seq to identify mediators of Alzheimer’s disease risk"

| | Sheet |
|---|---|
| 1 | Healthy |
| 2 | prAD |
| 3 | ABA |
| 4 | APO4 - 2 groups |

**This file contains supplementary tables as described below.**

| Description |
|---|
| ROSMAP dataset sex-specific differential expression genes analysis between the APOE group carreris vs. no-carriers |
| ROSMAP dataset sex-specific differential expression genes analysis between non-carriers and APOE4 carriers in prAD assigned individuals (prAD prob > 0.5) |
| ROSMAP dataset sex-specific differential expression genes analysis between non-carriers and APOE4 carriers in ABA assigned individuals (ABA prob > 0.5) |
| ROSMAP dataset differential expression genes analysis between non-carriers and APOE4 carriers (all individuals) |
