## Supplementary Table 3 for "DeepDynamics resolves cell-subtype and clinicopathological dynamics from bulk RNA-seq to identify mediators of Alzheimer’s disease risk"

**This file contains supplementary tables**

| | Sheet |
|---|---|
| 1 | ANOVA dynamics *APOE4* |
| 2 | ANOVA dynamics sex |
| 3 | ANOVA dynamics *APOE4* & sex |
| 4 | ANOVA pathways dynamics *APOE4* |

**as described below.**

| Description |
|---|
| ANOVA p-values on GAM models of dynamics, null vs. with *APOE4* factor |
| ANOVA p-values on GAM models of dynamics, null vs. with sex factor |
| ANOVA p-values on GAM models of dynamics, null vs. with *APOE4* factor for *APOE4* carriers and non-carriers separatley |
| ANOVA p-values on GAM models of pathways score (snRNA) dynamics, null vs. with *APOE4* factor |
