## Supplementary Table 2 for "DeepDynamics resolves cell-subtype and clinicopathological dynamics from bulk RNA-seq to identify mediators of Alzheimer’s disease risk"

**This file contains supplementary**

| | Sheet Name |
|---|---|
| 1 | DNN model predictions |
| 2 | Cell dynamics snuc |
| 3 | Cell dynamics bulk |
| 4 | Cell dynamics bulk e4 |
| 5 | Cell dynamics bulk sex |
| 6 | Cell dynamics e4+sex |
| 7 | snRNA pathways dynamics e4 |

**tables as described below.**

| Description |
|---|
| Pseudotime and trajectory probabilites predictions for each bulk RNA-seq sample and additional snRNA-seq |
| Dynamics of cell subpopulations and AD-associated traits from snRNA-seq data. |
| Cell subpopulations dynamics from bulk RNA-seq data. |
| Cell subpopulations dynamics from bulk RNA-seq data, starified by APOE4 genotype. |
| Cell subpopulations dynamics from bulk RNA-seq data, stratified by sex. |
| Cell subpopulations dynamics from bulk RNA-seq data, stratified by both APOE4 genotype and sex. |
| snRNA pathway dynamics startified by APOE4 genotype. |
