## Supplementary Table 1 for "DeepDynamics resolves cell-subtype and clinicopathological dynamics from bulk RNA-seq to identify mediators of Alzheimer’s disease risk"

**This file**

| Sheet |
|---|
| Metadata |
| Bulk supboulations abundance |

**contains supplementary tables as described below.**

| Description |
|---|
| Metadata that was used for the individuals in the study. |
| Cell subpopulations prevalances predicted by CelMod for each individual. |
